# MTHFR*677C>T produces distinct prodromal disease signatures in a mouse model of late-onset Alzheimer’s disease

**DOI:** 10.64898/2026.08.25.746973

**Authors:** Kevin P. Kotredes, Ravi S. Pandey, Alaina M. Reagan, Zeynep Sarica, Rita O’Rourke, Sarah Herrick, Avery Davis, Dylan Garceau, Michael Sasner, Gregory W. Carter, Gareth R. Howell

**Affiliations:** The Jackson Laboratory, 600 Main Street, Bar Harbor, ME, United States 04609; The Jackson Laboratory for Genomic Medicine, 10 Discovery Drive, Farmington, CT, United States 06032; Tufts University Graduate School of Biomedical Sciences, 136 Harrison Ave #813, Boston, MA, United States 02111; University of Maine, 5775 Stodder Hall, Orono, Maine, United States 04469

**Keywords:** Late-onset Alzheimer’s disease, LOAD2, Methylene tetrahydrofolate reductase, Mthfr, Mouse model, Neuropathology, Transcriptomics, Proteomics, Aging, Cerebrovascular deficits

## Abstract

**Background:** Late-onset Alzheimer’s disease (LOAD) comprises more than 95% of all AD cases. Transgenic, overexpression animal models have off target side effects, do not effectively produce the heterogeneity observed clinically in LOAD patients, and are therefore not best suited for preclinical therapeutic development. The Model Organism Development and Evaluation for Late-onset Alzheimer’s Disease (MODEL-AD) Consortium was established to develop novel mouse strains to model human-relevant genetic and environmental risk factors for LOAD. Methylenetetrahydrofolate reductase (MTHFR) is an enzyme in the folate/methionine pathway. Variants in the *MTHFR* gene, notably *677C>T,* are associated with ADRD, and we have previously shown the *Mthfr^677C>T^*mouse model phenocopies humans carrying the variant and develop cerebrovascular deficits.

**Methods:** To examine the contributions of *Mthfr^677C>T^*in the context of late-onset Alzheimer’s disease (LOAD), MODEL-AD created a novel mouse strain on the C57BL/6J (B6) background that was homozygous for *Mthfr^677C>T^*, in combination with humanized *Aβ*, *APOEe4*, and *Trem2*R47H* (*referred to as LOAD2.Mthfr^677C>T^)*. Mice were assessed over multiple ages for disease-relevant phenotypes. Regular behavior measurements and biometric samples were collected longitudinally to 24 months of age. Blood and brain tissue were collected for transcriptomics, proteomics, human disease correlation, and neuropathology.

**Results:** Despite lacking hallmark pathologies such as amyloid deposition and significant neuroinflammation, compared to LOAD2 controls, *LOAD2.Mthfr^677C>T^*mice showed transcriptional and proteomic signatures in the brain that relate to the cerebrovasculature, myelination, and synaptic biology, similar to those seen in human LOAD patients.

**Conclusions:** These data further support the use of the LOAD2.Mthfr*^677C>T^*mouse model to study aspects of ADRD such as cerebrovascular compromise.

## BACKGROUND

Alzheimer’s disease (AD) pathology was first described in dementia patients as exhibiting senile plaque and neurofibrillary tangle aggregates in the cerebral cortex [1]. These foundational pathological hallmarks were the first evidence of the cellular processes that produce neuron loss and neurobehavioral phenotypes [2]. The evolution of biomedical technologies has since allowed for improved description and expanded classification of senile dementia. In addition to AD, it is widely accepted that there are other types of related dementias (RD), attributed to either vascular deficits, other proteinaceous deposits, or brain atrophy [3–9]. Increased risk for ADRDs comes from specific genetic alterations that have been unveiled by next-generation-sequencing and gene editing technologies [10, 11]. As the availability of these human disease reference datasets grows, better defined genetic risk factors in combination with environmental risk factors are emerging, correlating to the prevalence or magnitude of disease burden [12, 13]. With this information, the field has grown to appreciate that neurodegenerative disorders are linked to nearly all aspects of intra-and extra-cellular homeostatic dynamics, including metabolism, immunity, cell cycle regulation, and DNA repair and regulation [14–17].

Unlike other age-related diseases that have seen precipitous declines in the rates of incidence and severity - thanks to advances in prevention, diagnostics, and therapeutics – AD and ADRD continues to be more prevalent [18–20]. It is estimated that the risk of diagnosis doubles every five years following 65 years of age [21]. In the absence of impactful interventions, increases in life expectancy have outpaced preventative habits to continually place more burden on healthcare. To counteract the growth of the AD patient population, better efforts must be made to improve therapeutics. This begins at the preclinical stage, where animal studies elucidate disease processes at the subcellular, cellular, and tissue level. These insights then inform appropriate interventions that may be applied to human medicine. Therefore, novel preclinical approaches and better disease models are needed to supplement the legacy toolkits developed decades ago. To overcome this gap in knowledge, the MODEL-AD (Model Organism Development and Evaluation for Late-Onset Alzheimer’s Disease) consortium was established to identify novel, human-relevant genetic risk factors and engineer mouse strains to better model the complexity of human dementias, including late-onset AD (LOAD) [22, 23].

We have previously shown that the LOAD2 mouse model – that is triple homozygous for humanized Aβ, *APOE4*, and *Trem2^R47H^*– showed signatures of synaptic dysfunction and cognitive deficits, in the absence of amyloid and tau pathology [24]. Here we now described the creation and phenotyping of the LOAD2.*Mthfr^677C>T^* mouse strain, developed to investigate how the *Mthfr^677C>T^* genetic risk factor may synergize with other LOAD risk factors to influence processes relevant to LOAD. Methylenetetrahydrofolate reductase (MTHFR) is a critical enzyme in one-carbon metabolism. MTHFR catalyzes the reduction of 5, 10-methylenetetrahydrofolate to 5-methyltetrahydrofolate, the primary form of circulating folate and the carbon donor for homocysteine re-methylation to methionine. The *MTHFR*\*677C>T polymorphism is a common variant [25] that has been associated with increased risk for several disorders, including stroke, coronary disease and hypertension as well as with Alzheimer’s disease [26–28] and vascular dementia [29, 30]. This gene variant encodes an alanine to valine substitution at position 222 (A222V) that impairs enzyme function and leads to elevated plasma homocysteine, a risk factor for vascular inflammation [31]. Because 20-40% of the general population carries at least one copy of the C>T variant [32], understanding its contribution to vascular risk is broadly relevant.

Our group previously generated and characterized the B6.*Mthfr^677C>T^*knock-in mouse model [33], demonstrating that mice have reduced MTHFR activity and increased plasma homocysteine in a clinically relevant manner [33]. Mice also display cerebrovascular phenotypes, including reduced vascular density in frontal cortex and reduced blood perfusion by PET, suggesting that the variant contributes to cerebrovascular dysfunction [33]. We also showed *Mthfr^677C>T^* in the context of *APOE4* and *Trem2^R47H^* (LOAD1) mice modeled molecular signatures of clinical LOAD [34], that were not increased by high fat diet [35]. Collectively, these studies support incorporating the *Mthfr^677C>T^*variant into mouse models to understand the genetic complexity of LOAD. Here, we now show that, despite lacking hallmark pathologies such as amyloid and tau, compared to LOAD2 or B6 controls, LOAD2.*Mthfr^677C>T^* mice show strong age-and sex-dependent transcriptomic and proteomic signatures relevant to human LOAD, including in processes relevant to cerebrovascular, myelination, and synapse biology. This new strain adds to the toolbox of mouse models for LOAD, created by MODEL-AD, for more precise preclinical testing of novel therapeutic approaches to treat LOAD.

## METHODS

*Mouse strain development:* The LOAD2.*Mthfr^677C>T^*mouse strain (JAX stock 036242), as well as background, control LOAD2 (JAX stock 030670) mice, were designed to model human LOAD by expressing risk alleles strongly linked to increased disease incidence and severity [36]. Mouse *Apoe* transcripts were mutated to express a humanized sequence coding for the human isoform ‘E4’ of *APOE*, including exons 2, 3 and 4 (and some 3’ UTR sequence) [34, 37]. The *Trem2* allele was generated by inserting a missense point mutation R47H in exon 2 [34, 37, 38] and *App* was humanized with mutations at G601R, F606Y, and R609H which correspond to amino acid positions 676, 681, 684 in the human *APP* locus encoding for amyloid beta (Abeta) region 1-42 [24]. Detailed descriptions of model generation and phenotyping of baseline alleles expressed in the ‘LOAD2’ mouse strain were published previously [24]. Mouse methylenetetrahydrofolate reductase (*Mthfr)* was targeted by CRISPR/Cas9 to introduce a point mutation of A262V that replicates the polymorphism (*Mthfr*^677C>T^) associated with increased risk of Alzheimer’s disease in human genome-wide association studies [34, 39, 40].

For additional strain information including genotyping, husbandry, development, and availability, visit The Jackson Laboratory Mouse Search: e.g. https://www.jax.org/strain/036242 *Cohorts:* Six cohorts of LOAD2.*Mthfr^677C>T^* mice and controls were created at The Jackson Laboratory to evaluate LOAD-relevant phenotypes:

|  | Genotypes | Sex | n= |
| --- | --- | --- | --- |
| 4-month | B6 | M/F | 6/6 |
|  | LOAD2 | M/F | 12/12 |
|  | LOAD2.*Mthfr*<sup>677C>T</sup> | M/F | 12/12 |
| 12-month | B6 | M/F | 6/6 |
|  | LOAD2 | M/F | 12/12 |
|  | LOAD2.*Mthfr*<sup>677C>T</sup> | M/F | 12/12 |
| 18-month | B6 | M/F | 6/6 |
|  | LOAD2 | M/F | 12/12 |
|  | LOAD2.*Mthfr*<sup>677C>T</sup> | M/F | 11/12 |
| 24-month | B6 | M/F | 6/6 |
|  | LOAD2 | M/F | 12/12 |
|  | LOAD2.*Mthfr*<sup>677C>T</sup> | M/F | 9/11 |

To produce experimental cohorts, LOAD2.*Mthfr^677C>T^*heterozygote (B6J.*APOE^E4/E4^*.*Trem2^R47H/R47H^*.*App^hAb/hAb^.Mthfr^677C>T/+^*) mice were intercrossed to create B6J.*APOE^E4/E4^*.*Trem2^R47H/R47H^*.*App*^hAb/hAb^ (LOAD2, triple homozygous) and B6J.*APOE^E4/E4^*.*Trem2^R47H/R47H^*.*App^hAb/hAb^.Mthfr^677C>T/677C^*(LOAD2.*Mthfr^677C>T^*, quadruple homozygous) genotype littermate mice. In addition to these animals, C57BL/6J (B6) control animals were aged and tested in parallel with experimental LOAD2.*Mthfr^677C>T^*cohorts. In appreciation of sexual dimorphism observed in human aging and disease, male and female mice were established for a combination of longitudinal and cross-sectional phenotyping at 4-, 12-, 18, and 24 months. The 18-month group was assessed for biometrics and plasma biomarkers at 4-, 8-, 12-and 18 months of age.

### Housing

All procedures were approved by The Jackson Laboratory Institutional Animal Care and Use Committee (IACUC). Mice were bred and maintained in a 12/12-hour light/dark cycle, consisting of 12 hour-ON (6:00 am-6:00 pm), followed by 12 hour-OFF. Room temperatures are maintained at 18–24°C (65–75°F) with 40–60% humidity. All mice were housed in positive, individually ventilated cages (PIV). Standard autoclaved 6% fat diet (Purina Lab Diet 5K52) was available to the mice *ad-lib*, as was regular drinking water with acidity regulated from pH 2.5–3.0. At 2 months of age, each experimental group was randomized. The animals were ear-punched for identification purposes and subsequently microchipped at the tail base using a p-chip system (PharmaSeq). All behavioral characterization was conducted in the Neurobehavioral Phenotyping core (NBP) in the Center for Biometric Analysis (CBA) at The Jackson Laboratory. Briefly, mice were relocated from the housing room, in which they were reared, to the adjacent NBP facility. Mice were individually housed at minimum 5 days prior to behavioral testing. The dedicated CBA housing room consists of PIV caging with temperature controlled at a setting of 22 ± 1°C (72 ± 2°F) and humidity at 50 ± 20%. The testing facility was on a 12:12 L:D schedule (lights on at 6:00 am) with all testing performed during the light cycle (typically between 7:00 am and 5:00 pm). All subjects were randomized and counterbalanced for testing order across multiples of instrumentation and time of day for each test day, with a simplified testing ID number (e.g., #1-100), with all technicians blinded to genotype (e.g., coded as A, B, C, etc.). The blind was maintained throughout testing and until after the data were analyzed with no subjects or data excluded based on any mathematical outliers.

### Behavior

#### Frailty assay

The frailty assessment was conducted as previously published [24, 41] and is used to assess the presence of a spectrum of aging related characteristics in mice. Briefly, test subjects were transported to the testing room, body weights were recorded, and tails were labeled with the subject ID number (e.g. 1-50) using a non-toxic marker. Mice were then left undisturbed for a minimum of 60 minutes to acclimate to the procedure room. Following acclimation, subjects were individually evaluated for the absence or presence of 27 characteristic traits and reflexes and scored a 0, 0.5, or 1 (based on presence/absence, and severity) for each assessment by a trained observer, blind to genotype/age, and included the following assessments: alopecia; loss of fur color; dermatitis/skin lesions; loss of whiskers; coat condition; piloerection; cataracts; eye discharge/swelling; microphthalmia; nasal discharge; rectal prolapse; vaginal/uterine/penile; diarrhea; vestibular disturbance; vision loss assessed by visual placing upon subject being lowered to a grid; menace reflex; tail stiffening; impaired gait during free walking; tremor; tumors; distended abdomen; kyphosis; body condition; breathing rate/depth; malocclusions; righting reflex; hindlimb clasping. The frailty index score was calculated as the cumulative score of all measures with a maximum score of 27. Frailty index score fold change is plotted following normalization across all three genotypes to set 4-month measurements, for all animals, to equal 1. Core body temperature was recorded just prior to the conclusion of the frailty assessment via a glycerol lubricated thermistor rectal probe (Braintree Scientific product# RET 3; measuring 3/4” in length, .028” diameter, .065” tip) inserted ∼2 cm into the rectum of a manually restrained mouse for approximately 10 seconds. Temperature was recorded to the nearest 0.1°C (#TH5 Thermalert digital thermometer, Braintree Scientific, Braintree, MA).

#### Open field test

Versamax Open Field Arenas (40 cm x 40 cm x 40 cm; Omnitech Electronics, OH USA) were used for this test. Arenas were housed within sound attenuated chambers with lighting in the testing room and arenas consistent with the housing room (∼500 lux). Mice were placed individually into the center of the arena and infrared beams recorded distance traveled (cm), vertical activity, and perimeter/center time. Data were collected in 5-minute time bins for a duration of 60 minutes.

#### Spontaneous alternation

Mice were acclimated to the testing room under ambient lighting conditions (∼20-50 lux). For this test, a clear polycarbonate Y-maze (in-house fabricated; arm dimensions 33.65 cm length, 6 cm width, 15 cm height) with no intended visual cues, was placed on top of an infrared reflecting background (Noldus, The Netherlands), surrounded by a black floor-to ceiling curtain to minimize extra-maze visual cues Mice were placed midway of the start arm (A), facing the center of the ‘Y’ for an 8-minute test period in which the center point of the mouse was tracked for the sequence of entries into each of the three arms (labeled A, B and C) which were recorded via a ceiling-mounted infrared camera integrated with behavioral tracking software (Noldus Ethovision XT). Percent spontaneous alternation is calculated as the number of correct triads (entries into each of the three different arms of the maze in a sequence of three without returning to a previously visited arm, e.g. ABC, CAB, BCA) relative to the number of alteration opportunities (number of total triads). For example, in a sequence of the following entries: ACBACCBACBCA; the triads are denoted as follows: ACB, CBA, BAC, ACC, CCB, CBA, BAC, ACB, CBC, BCA. Therefore, the analyzed data would have identified 7 correct sequences out of a total of 10 alternation opportunities = 70% correct alternations.

#### Rotarod Test for Motor Coordination

An accelerating Rotarod (Ugo-Basile; model 47600) is used for this test. Lighting in the testing room is consistent with the housing room (∼500 lux). The trial begins with mice being placed on the rotating rod (4 rpm), which accelerates up to 40 rpm over the course of 300 seconds. Each mouse is subjected to 3 consecutive trials with an ∼1-minute inter-trial interval to allow cleaning of the rod between trials. Latency to fall (seconds) is measured. Subjects that fall upon initial placement on the rod, before acceleration begins, are scored as 0 seconds for that trial.

#### Blood plasma collection

Blood was collected longitudinally from non-fasted mice via cheek puncture. For terminal timepoint collections, non-fasted blood was collected at harvest. The chest cavity of an anesthetized mouse was opened to reveal the heart. With a 25G EDTA-coated needle, blood was extracted by cardiac puncture from the left ventricle prior to PBS perfusion. Approximately 500 µL of whole blood was then placed in a MAP-K2 EDTA Microcontainer (BD, Franklin Lakes, NJ) on ice. Microcontainers were then spun at 5,000 rpm (4,388 x g) in a pre-chilled ultracentrifuge at 4°C for 15 minutes. Without disturbing red blood cell fraction, plasma supernatant was pipetted with p200 tip into chilled cryovial. The cryovial was then snap-frozen immediately with dry ice for at least 10 minutes before placing in −80°C for storage.

### Brain tissue collection

#### Animal anesthetization and perfusion

Upon arrival at the terminal endpoint for each mouse cohort, individual animals were weighed prior to intraperitoneal administration of tribromoethanol (1 mg/kg). Deep anesthetization via toe pinch is confirmed. If desired, prior to perfusion blood and CSF samples are collected. Next, an incision along the ventral midline is made to expose the thorax and abdomen, followed by removal of the lateral borders of the diaphragm and ribcage to reveal the heart. To perfuse the animal, a small cut was placed in the right atrium to relieve pressure from the vascular system before perfusing the animal transcardially with 1X PBS via injection into the left ventricle. Completion of perfusion and clearance of the vascular system was indicated by a blanching of the liver.

#### Histological sample collection

During harvest, whole mouse brains were removed and weighed. Using a brain matrix, left and right hemispheres were separated along the midsagittal plane. Right brains were dissected and snap frozen for preservation of RNA for gene expression studies. The left hemisphere was placed in 5 mL of 4% PFA at 4°C overnight, then moved to 10 mL of 15% sucrose at 4°C overnight or until brain sinks to bottom of the tube, before finally being incubated in 10 mL of 30% sucrose at 4°C overnight or until brain sinks to bottom of the tube. The left hemisphere was then snap frozen and stored at −80°C until sectioned.

#### ELISAs

Right hemisphere brain samples and plasma were assayed in duplicate using the V-PLEX MesoScale Discovery (MSD) Mouse Proinflammatory Panel I, a highly sensitive multiplex enzyme-linked immunosorbent assay (ELISA) for quantitatively measuring cytokines including interferon γ (IFN-γ), interleukin (IL)-1β(beta), IL-2, IL-4, IL-5, IL-6, IL-10, IL-12p70, KC/GRO, and tumor necrosis factor α (TNFα [alpha]) (K15048D, MesoScale Discovery, Gaithersburg, MD, USA). MSD human (amyloid) Aβ Peptide Panel I was employed, similarly, to quantify plasma Aβ1-38, 1-40, and 1-42 (K15200E). Neurofilament light chain (Nf-L) levels in plasma samples were quantified using Simple Plex Mouse NF-L Cartridges on the Ella Automated ELISA System (Bio-Techne). Plasma was analyzed by Siemens Advia 120 (Germany) for levels of homocysteine, glucose, total cholesterol, LDL (low-density lipoproteins), HDL (high-density lipoproteins), triglycerides, and NEFA (non-essential fatty acids).

#### Immunohistochemistry

Following PFA and sucrose gradients, left brain hemispheres were sectioned on a Thermo Scientific HM430 sliding microtome at 25 μm thickness. Coronal brain tissue sections, oriented to capture the cortex and hippocampus at approximately: Bregma: −2.75 mm and Interaural 1.05 mm, were collected in 1X PBS. Using the hippocampus as a landmark all sections with CA1, 2, and 3 are collected). Additional images and subsequent cell densities of the dorsal subiculum (and nearby cortex [primary visual area] region) were obtained around bregma target −3.60 mm. One hundred coronal slices are collected per brain – encapsulating about 2.5 mm of the horizontal plane. Floating sections were then blocked prior to immunohistochemical staining and mounting. After blocking slides with 10% normal donkey serum or normal goat serum diluted in 1X PBS + 0.5% Triton wash buffer, floating sections were incubated with antibodies selected to visualize neurons (NeuN; ab104225, 1:1000, Abcam), astrocytes (GFAP; AP31806PU-N, 1:1000, Origene), microglia (Iba1; ab178847, 1:500, Abcam), plaques (X-34 or 6E10; 803001, 1:1000, Biolegend), and nucleus (DAPI). Ten sections, collected in equal representation throughout the brain, are arranged on each microscope slide. Images were acquired at 20X using the Leica Aperio Versa (Germany) slide scanner or Leica Thunder microscope (Germany). Regions of the cortex, subiculum, and hippocampus (∼650 µM x 650 µM) were processed using Imaris (Chicago, IL, USA) software to quantify cell counts, fluorescence intensity, and surface area ratios. Counts/density adjusted to positive cells/mm^2^.

### Transcriptomics

#### Sample extraction

Total RNA was isolated from tissue using the NucleoMag RNA Kit (Macherey-Nagel) and the KingFisher Flex purification system (ThermoFisher). Tissues were homogenized in MR1 buffer (Macherey-Nagel) using a Bead Ruptor Elite (Omni International). RNA isolation was performed according to the manufacturer’s protocol. RNA concentration and quality were assessed using the Nanodrop 8000 spectrophotometer (Thermo Scientific) and the RNA ScreenTape Assay (Agilent Technologies).

#### Library Preparation

Stranded libraries were constructed using the KAPA mRNA HyperPrep Kit (Roche Sequencing and Life Science), according to the manufacturer’s protocol. Briefly, the protocol entails isolation of polyA containing mRNA using oligo-dT magnetic beads, RNA fragmentation, first and second strand cDNA synthesis, ligation of Illumina-specific adapters containing a unique barcode sequence for each library, and PCR amplification. The quality and concentration of the libraries were assessed using the D5000 ScreenTape (Agilent Technologies) and Qubit dsDNA HS Assay (ThermoFisher), respectively, according to the manufacturers’ instructions.

#### Sequencing

Libraries were pooled and sequenced by the Genome Technologies core facility at The Jackson Laboratory. All samples were sequenced 150 bp paired-end on an Illumina NovaSeq X Plus using the 10B Reagent Kit (Illumina), targeting 30 million read pairs per sample. Once the data was received the samples were concatenated to have a single file for paired-end analysis.

#### Data processing

RNA-Seq data were processed using nf-core/rnaseq pipeline [https://doi.org/10.5281/zenodo.1400710]. Briefly, reads were aligned to the reference mouse genome (version GRCm38.p6) using STAR [42] and gene expression was quantified with RSEM [43]. To measure human APOE gene expression, we created a custom mouse reference genome by concatenating human APOE gene sequence (human chromosome 19:44905754-44909393; build GRCh38.p10) into the mouse genome (GRCm38.p6) as a separate chromosome (referred as chromosome 21 in chimeric mouse genome). Subsequently, we added a gene annotation for the human APOE gene into the mouse gene annotation file. Genotypes were validated by using samtools mpileup to count nucleotide occurrences at relevant positions and compare the number of reference vs alternate alleles. Sex was validated by comparing expression of the sex-specific genes Xist (ENSMUSG00000086503), Eif2s3y (ENSMUSG00000069049), and Ddx3y (ENSMUSG00000069045)

### Proteomics

#### Tissue Homogenization and Protein Digestion

Samples were homogenized in 8 M urea lysis buffer (8 M urea, 10 mM Tris, 100 mM NaH2PO4, pH 8.5) with HALT protease and phosphatase inhibitor cocktail (ThermoFisher) using a Bullet Blender (NextAdvance). Each Rino sample tube (NextAdvance) was supplemented with ∼100 μL of stainless-steel beads (0.9 to 2.0 mm blend, NextAdvance) and 300 μL of lysis buffer. Tissues were added immediately after excision and homogenized with bullet blender at 4 °C with 2 full 5-minute cycles. The lysates were transferred to new Eppendorf Lobind tubes and sonicated for 3 cycles consisting of 5 seconds of active sonication at 30% amplitude, followed by 15 seconds on ice. Samples were then centrifuged for 5 minutes at 15,000 x g and the supernatant transferred to a new tube. Protein concentration was determined by bicinchoninic acid (BCA) assay (Pierce). For protein digestion, 100 μg of each sample was aliquoted and volumes normalized with additional lysis buffer. Samples were reduced with 5 mM dithiothreitol (DTT) at room temperature for 30 minutes, followed by 10 mM iodoacetamide (IAA) alkylation in the dark for another 30 minutes. Lysyl endopeptidase (Wako) at 1:25 (w/w) was added, and digestion allowed to proceed overnight. Samples were then 7-fold diluted with 50 mM ammonium bicarbonate. Trypsin (Promega) was then added at 1:25 (w/w) and digestion proceeded overnight. The peptide solutions were acidified to a final concentration of 1% (vol/vol) formic acid (FA) and 0.1% (vol/vol) trifluoroacetic acid (TFA) and desalted with a 30 mg HLB column (Oasis). Each HLB column was first rinsed with 1 mL of methanol, washed with 1 mL 50% (vol/vol) acetonitrile (ACN), and equilibrated with 2×1 mL 0.1% (vol/vol) TFA. The samples were then loaded onto the column and washed with 2×1 mL 0.1% (vol/vol) TFA. Elution was performed with 2 volumes of 0.5 mL 50% (vol/vol) ACN.

#### Isobaric Tandem Mass Tag (TMT) Peptide Labeling

Each sample (containing 20 μg of peptides) was re-suspended in 100 mM TEAB buffer (100 μL). The TMT labeling reagents (5 mg) were equilibrated to room temperature, and anhydrous ACN (256 μL) was added to each reagent channel. Each channel was gently vortexed for 5 minutes, and then 41 μL from each TMT channel was transferred to the peptide solutions and allowed to incubate for 1 hour at room temperature. The reaction was quenched with 5% (vol/vol) hydroxylamine (8 μL) (Pierce). All channels were then combined and dried by SpeedVac (LabConco) to approximately 150 μL and diluted with 1 mL of 0.1% (vol/vol) TFA, then acidified to a final concentration of 1% (vol/vol) FA and 0.1% (vol/vol) TFA. Labeled peptides were desalted with a 200 mg C18 Sep-Pak column (Waters). Each Sep-Pak column was activated with 3 mL of methanol, washed with 3 mL of 50% (vol/vol) ACN, and equilibrated with 2×3 mL of 0.1% TFA. The samples were then loaded and each column was washed with 2×3 mL 0.1% (vol/vol) TFA, followed by 2 mL of 1% (vol/vol) FA. Elution was performed with 2 volumes of 1.5 mL 50% (vol/vol) ACN. The eluates were then dried to completeness using a SpeedVac.

#### High-pH Off-line Fractionation

Dried samples were re-suspended in high pH loading buffer (0.07% vol/vol NH4OH, 0.045% vol/vol FA, 2% vol/vol ACN) and loaded onto a Water’s BEH 1.7 um 2.1mm by 150mm. An Thermo Vanquish was used to carry out the fractionation. Solvent A consisted of 0.0175% (vol/vol) NH4OH, 0.01125% (vol/vol) FA, and 2% (vol/vol) ACN; solvent B consisted of 0.0175% (vol/vol) NH4OH, 0.01125% (vol/vol) FA, and 90% (vol/vol) ACN. The sample elution was performed over a 25-minute gradient with a flow rate of 0.6 mL/min. A total of 192 individual equal volume fractions were collected across the gradient and subsequently pooled by concatenation into 96 fractions and dried to completeness using a SpeedVac.

#### Liquid Chromatography Mass Spectrometry

All fractions were resuspended in an equal volume of loading buffer (0.1% FA, 0.03% TFA, 1% ACN) and analyzed by liquid chromatography coupled to tandem mass spectrometry. Peptide eluents were separated on custom made fused silica column (15 cm × 150 μM internal diameter (ID) packed with Dr. Maisch 1.5um C18 resin) by a Vanquish Neo (ThermoFisher Scientific). Buffer A was water with 0.1% (vol/vol) formic acid, and buffer B was 80% (vol/vol) acetonitrile in water with 0.1% (vol/vol) formic acid. Elution was performed over a 20-minute gradient. The gradient was from 1% to 99% solvent B. Peptides were monitored on a Exploris 480 mass spectrometer (ThermoFisher Scientific) fitted with a high-field asymmetric waveform ion mobility spectrometry (FAIMS Pro) ion mobility source (ThermoFisher Scientific). Two compensation voltages (CV) of −45 and −65 were chosen for the FAIMS. Each cycle consisted of one full scan (MS1) was performed with an m/z range of 410-1600 at 120,000 resolution at standard settings and as many tandem (MS/MS) scans in 1.5 seconds. The higher energy collision-induced dissociation (HCD) tandem scans were collected at 32% collision energy with an isolation of 0.7 m/z, a resolution of 30,000* with TurboTMT on, an AGC setting of 250% normalized agc target, and a maximum injection time set to automatic. Dynamic exclusion was set to exclude previously sequenced peaks for 20 seconds within a 10-ppm isolation window.

#### Database Search

The raw files (1248 in total across 13 TMT batches) were searched using FragPipe (version 21.1). The FragPipe pipeline relies on MSFragger (version 4.0 [44, 45]) for peptide identification and Philosopher (version 5.1.0 [46]) for FDR filtering and downstream processing. The mouse protein database used contains canonical isoforms from Uniprot/Swissprot as of 02/2023 as well as the sequence for the human APOE protein along with specific peptides for the APOE2 and APOE4 alleles (CLAVYQAGAREGAER and LGADMEDVRGR respectively). The workflow used in FragPipe followed default TMT-16 plex parameters, used for both TMT-16 and TMT-18 experimental design. Briefly, precursor mass tolerance was −20 to 20 ppm, fragment mass tolerance of 20 ppm, mass calibration and parameter optimization were selected, and isotope error was set to −1/0/1/2/3. Enzyme specificity was set to strict-trypsin and up to two missing trypsin cleavages were allowed. Peptide length was allowed to range from 7 to 50 and peptide mass from either 200 to 5,000 Da. Variable modifications that were allowed in our search included: oxidation on methionine, N-terminal acetylation on protein, and N-terminal acetylation on peptide, with a maximum of 3 variable modifications per peptide. Peptide Spectral Matches were validated using Percolator [47]. The false discovery rate (FDR) threshold was set to 1% and protein and peptide abundances were quantified using Philosopher for downstream analysis.

#### Protein Quantitation and Quality Control

The protein abundances are normalized by scaling total protein signal within each channel for each specific case sample to the maximum channel-specific total signal. We then used a tunable median polish approach, TAMPOR, to remove technical batch variance in the proteomic data, as previously described [48]. TAMPOR is utilized to remove intra-batch and inter-batch variance while preserving meaningful biological variance in protein abundance values, normalizing to the median of selected intra-batch samples. This approach is robust to outliers and columns with up to 50% values missing. If a protein had more than 50% samples with missing values, it was removed from the matrix. No imputation of missing values was performed for any cohort. For the current data, TAMPOR leverages the median protein abundance from the pooled Global Internal Standard (GIS) TMT channels as the denominators in both factors to normalize sample-specific protein abundances across the 13 batches of samples.

#### Differential gene and protein expression analysis

Differentially expressed genes in mouse models were identified using the R Bioconductor package DESeq2 (v1.16.1) [49]. We used the Benjamini-Hochberg corrected p-values with a significance threshold of 0.05 to identify differentially expressed genes. We performed differential protein expression analysis for each mouse model compared to age and sex-matched control mice using one-way ANOVA followed by post hoc correction using Tukey HSD test to match methods used in analogous human studies [50].

#### Linear regression Analyses

We determined the effects of each factor (sex and genetic variants) by fitting a multiple regression model using the lm function in R as [51]:

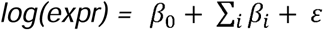

The sum is over sex (male), and all genetic variants *(LOAD2, Mthfr*677C>T)* used in this study. The expr represents gene expression measured by RNA-Seq transcript per million. In this formulation, B6 was used as the control for the LOAD2 mouse models, whereas LOAD2 served as controls for LOAD2.*Mthfr^677C>T^* models to estimate the effects of individual variants. Separate models were run for each age cohort.

#### Weighted gene co-expression network analysis of the mouse transcriptome

Weighted gene co-expression network analysis (WGCNA) [52] was performed to identify modules (clusters) of correlated genes using log TPM normalized expression values. We used the step-by-step construction approach for network construction and module identification [53]. The default unsigned network type was used, and a soft thresholding power of 5 was chosen to meet the scale-free topology criterion in the pickSoftThreshold function. We set the minimum modules size as 30, and merged modules whose correlation coefficient were greater than 0.75 (mergeCutHeight = 0.25). Each module is summarized by the module eigengene (ME), defined as first principal component of the gene expression profiles of each module. Further we computed the Pearson correlation coefficient of modules to genotype, age, sex, behavioral data, cytokines levels, and cell count data.

#### Functional enrichment analysis

Functional enrichment analyses were performed using the R Bioconductor package clusterProfiler [54], applying the enrichKEGG function with the org.Mm.eg.db annotation database to evaluate KEGG pathways. The significance threshold for all enrichment analyses was set at 0.05 using Benjamini–Hochberg adjusted *p*-values.

#### Human AMP-AD Gene Co-expression Modules

Wan et al. (2020) [55] discovered 30 human brain co-expression modules based on the meta-analysis of differential gene expression from seven distinct regions in postmortem samples obtained from three independent LOAD cohorts [56–58]. These modules were grouped into five distinct consensus clusters representing shared AD-related changes across studies and brain regions. Reactome pathway enrichment analysis was applied to annotate each cluster with distinct biological themes, ranking pathways by Bonferroni corrected *p*-values to ensure statistical rigor. Data for 30 human brain co-expression modules from the Accelerating Medicines Partnership for Alzheimer’s Disease (AMP-AD) studies were obtained from the Synapse data repository (https://www.synapse.org/#!Synapse:syn11932957/tables/; SynapseID: syn11932957). The log2 fold-change values for genes within these modules were originally computed from sex-regressed residualized counts. To enable direct comparisons between gene expression changes in female mice and female AD cases, and between male mice and male AD cases, we replaced these sex-regressed log2 fold-change values with sex-specific log2 fold-change values within each gene module. We obtained case–control log2 fold-change values for all quantified genes separately for males and females from the Synapse data repository (https://www.synapse.org/Synapse:syn30821563) and generated distinct male-specific and female-specific datasets for the 30 human brain co-expression modules.

#### Human LOAD Subtypes

Milind et al. (2020) [59] integrated post-mortem brain gene co-expression data from the frontal cortex, temporal cortex, and hippocampus due to their relevance to LOAD neuropathology across independent human cohorts (ROS/MAP, Mount Sinai Brain Bank, and Mayo Clinic) [56–58] and stratified patients into different molecular subtypes based on gene co-expression profiles using iterative WGCNA. Two distinct LOAD subtypes were identified in the ROSMAP cohort, three LOAD subtypes were identified in the Mayo cohort, and two distinct LOAD subtypes were identified in the MSBB cohort. Similar subtype results were observed in each cohort, with LOAD subtypes found to primarily differ in their inflammatory response based on differential expression analysis [59].

#### AD-related protein co-expression modules from human postmortem brain tissue

Human protein co-expression modules were obtained from previously published protein expression profiles from human dorsolateral prefrontal frontal cortex [50]. Briefly, Johnson et al. analyzed more than 500 dorsal prefrontal cortex (DLPFC) tissues from control, asymptomatic AD (AsymAD), and AD brains and generated a deep TMT AD protein network using WGCNA. This network consists of 44 modules of proteins related to one another by their co-expression across control and disease tissues. Johnson, et al. functionally annotated these modules using Gene Ontology analysis of its constituent proteins and assigned cell types to each module, and measured correlations of each summary module eigenprotein to neuropathological or cognitive traits present in the cohorts [50]. We obtained case-control LogFC values previously adjusted for sex for each quantified protein and membership in protein co-expression modules from the AD Knowledge Portal (https://www.synapse.org/#!Synapse:syn25453861).

#### Mouse-human correlation analysis

We assessed the similarity between mouse and human disease-related expression changes by calculating Pearson correlations between changes in expression (log2 fold change) in human AD cases versus controls and the changes in expression (log2 fold change) in each mouse model (LOAD2 vs. B6, LOAD2.*Mthfr^677C>T^* vs. B6, and LOAD2.*Mthfr^677C>T^* vs. LOAD2), stratified by age and sex. Correlations were computed across the set of orthologous genes in each AMP-AD module (and orthologous proteins within corresponding proteomics modules for proteomics data) [36, 51] using cor.test function in R as:

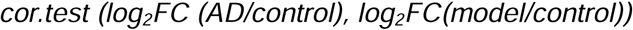

Additionally, we assessed correlation between human AD log2 fold-change values and the estimated effect of the *Mthfr^677C>T^* variant (β) from the linear model for mouse orthologs genes. Correlation coefficients were computed using cor.test function built in R as:

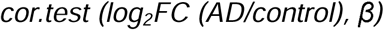

The cor.test function returns both the correlation coefficient and the significance level (p-value) of the correlation.

To compare mouse models with human LOAD subtypes, we computed the Pearson correlations between log2 fold-change values in each subtype versus controls and those in each mouse model, stratified by age and sex:

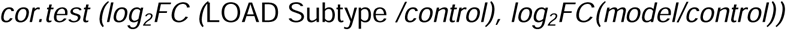

We plotted the correlation results using the ggplot2 package in R. Framed circles were used to denote significant (p < 0.05) positive (blue) and negative (red) Pearson’s correlation coefficients. The color intensity and size of the circles were sized proportional to Pearson’s correlation coefficient.

#### AD biological domains

Cary et al. [60] developed 19 biological domains that capture the AD-associated endophenotypes and defined them using an exhaustive set of Gene Ontology (GO) terms, with the intent to keep each domain siloed in a biologically coherent fashion. Genes and proteins were grouped into biodomains based on annotation to a term within each biodomain. For each biological domain, the Pearson correlation was computed between the estimated effect of the *Mthfr^677C>T^* variant (β) on each gene or protein and the meta-analysis treatment effect for each orthologous gene or protein from the human dataset (https://www.synapse.org/#!Synapse:syn22758536/tables/).

#### Statistical Analysis

Statistical analyses, except those related to transcriptomics and proteomics datasets, were performed on GraphPad Prism version 11.0.2 (GraphPad Software, Boston, MA, USA). Data are presented as mean ± SEM. Comparisons involving two experimental factors (e.g., genotype and treatment) were analyzed using two-way ANOVA with Tukey’s multiple comparisons test. Statistical significance was defined as p < 0.05.

## RESULTS

### LOAD2.Mthfr^677C>T^ do not display cognitive deficits but do show anxiety-like behavior

Longitudinal cohorts consisting of male and female C57BL/6J (B6), LOAD2, and LOAD2.*Mthfr^677C>T^* mice were created, sampled, and analyzed in parallel. Initial phenotyping occurred when animals were 4 months old and extended throughout life until mice they reached 24 months. *In vivo* neurobehavioral phenotyping proceeded as described previously [24, 37], including a battery of assays implemented to collectively describe the health status of experimental mice.

At 4 months of age, cumulative frailty index scores were expectedly very low for all three genotypes (FIGURE 1A), as frailty measurements are designed to be principally affected by age. Compared to 4-month baselines, longitudinal cohorts displayed increased frailty scores by 12-, 18-, and through to 24 months of age, regardless of sex or genotype. At each of these aged timepoints, LOAD2 and LOAD2.*Mthfr^677C>T^*animals’ scores were greater than those with a B6 genotype. LOAD2. *Mthfr^677C>T^*females at 18 months of age show the highest scores, which may suggest an accelerated aging phenotype.

**FIGURE 1:**
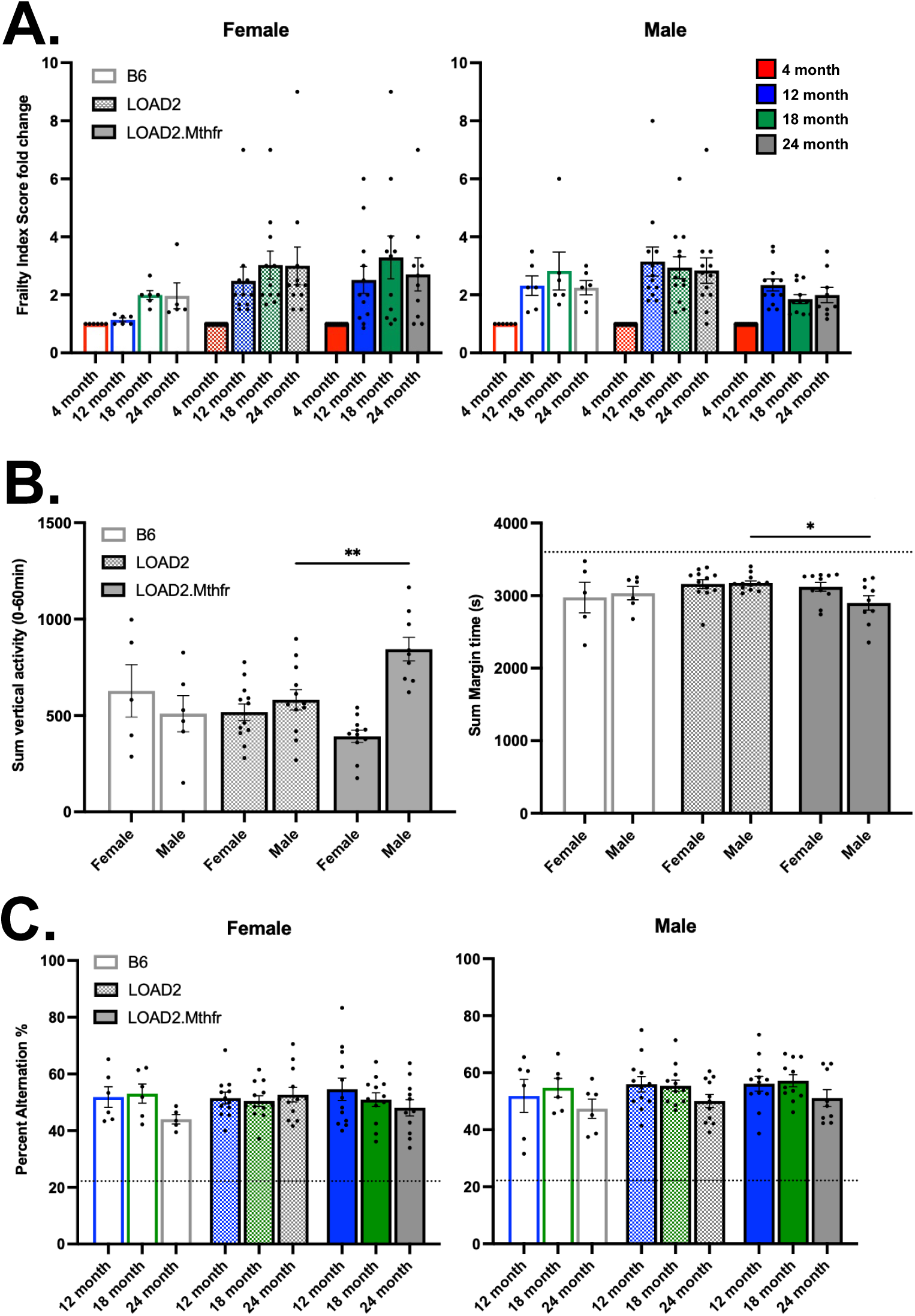
Longitudinal behavioral phenotyping of animal cohorts. C57BL/6J (B6), LOAD2, and LOAD2. *Mthfr^677C>T^*behavioral measurements. Frailty index assay performed longitudinally at 4-, 12-, 18-, and 24 months of age and changes plotted after normalized to 4-month timepoint for each genotype (A). Open field assay cross sectional measurements in 24-month-old animals for total number of vertical movements (rearing) and total margin time, in seconds, and over a 60-minute period (B). Spontaneous alternation performance by animal movements in Y-maze without visual cues, tested longitudinally at 12-, 18-, and 24 months of age. Random chance of completing full alternation equals 22.2% (C). Sample size n≥5. Significant statistical difference determined within sex groupings by ANOVA: *p<0.05, **p<0.01, ***p<0.001.

Analysis of animal behavior via open field assay showed that LOAD2.*Mthfr^677C>T^* males at 24 months exhibited hyperexcitability-related phenotypes: decreased margin time and increased vertical activity (rearing) (FIGURE 1B) compared to LOAD2 controls. Cognition was tracked by spontaneous alternation assay (Y-maze, without visual cues) through adulthood. All groups, irrespective of age, genotype or sex, were able to complete the task suggesting short term spatial working memory was intact (FIGURE 1C).

### LOAD2.Mthfr^677C>T^ and LOAD2 mice display sex-or genotype-specific changes in neurons, microglia and astrocytes

For evidence of neuropathological phenotypes, brains from aged (24 months) animals were probed for the presence of amyloid plaques and changes in neuronal and glial cell counts.

Representative images from the cortex, hippocampus, and subiculum were obtained by immunohistochemical analysis (SUPP FIGURE 1) and revealed that in the context of LOAD2, the *Mthfr^677C>T^* variant had little effect on total neuron cell counts (FIGURE 2A). Interestingly, there were pronounced sex-specific differences in NeuN-DAPI-double-positive cells in the subiculum, which may be relevant to the sex-specific difference we observe in anxiety-related behavior (FIGURE 1B).

**FIGURE 2:**
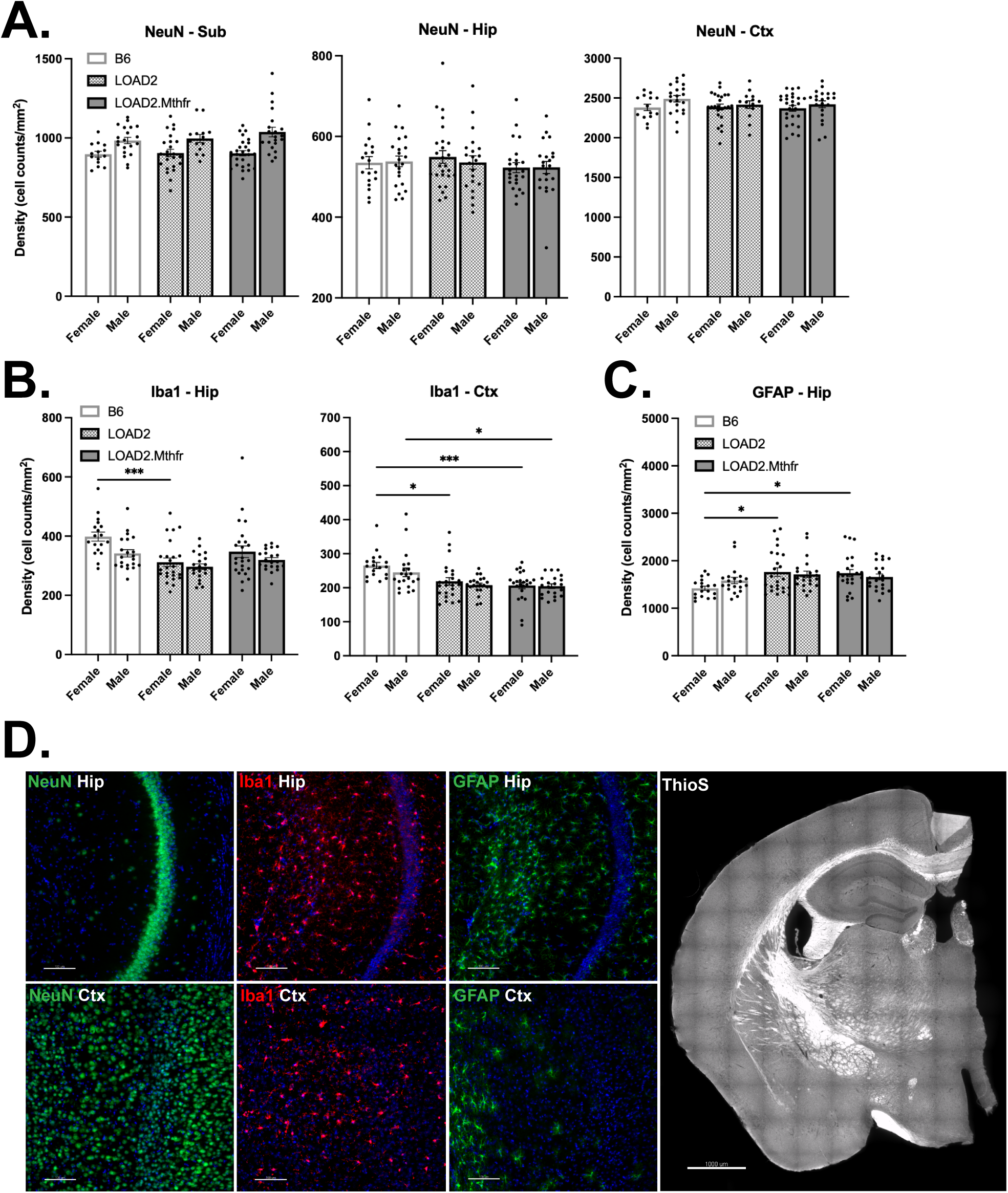
Neuropathological assessment of brains from aged LOAD mice expressing *Mthfr^677C>T^.* C57BL/6J (B6), LOAD2, and LOAD2.*Mthfr^677C>T^* 24-month-old cross-sectional cohort immunohistochemistry. Tissue sections 25 micron thick imaged at 20X magnification quantified per brain (bregma markers approximately −2.00 mm). Representative images obtained from the cortex (Ctx), hippocampus (Hip), and subiculum (Sub) and analyzed for cell counts of neurons (NeuN+) (A), microglia (Iba1+) (B), and astrocytes (GFAP+) (C); counterstained with DAPI+ (blue) (representative images) (D). Full hemisphere of ThioS staining depicting absence of amyloid plaques (D). Sample size n≥15. Significant statistical difference determined within sex groupings by ANOVA: *p<0.05, **p<0.01, ***p<0.001.

Assessment of microglia and astrocyte counts revealed sex-and genotype-specific differences. Microglial counts revealed genotype-driven changes in Iba1-DAPI-double positive cell populations (FIGURE 2B). When compared to female B6 mice, female LOAD2 mice show fewer Iba1-positive microglia in hippocampus and female LOAD2 and LOAD2.*Mthfr^677C>T^* mice show fewer in cortex. These differences were also observed at a young timepoint (SUPP FIGURE 2C), suggesting cell number differences may be determined early in life and are not exacerbated by aging in these models. Genotype-related changes were also observed in GFAP-positive astrocyte populations. In contrast, female LOAD2 and LOAD2.*Mthfr^677C>T^* mice had increased astrocyte counts and decreased density (FIGURE 2C; SUPP FIGURE 1D) compared to B6 controls in the hippocampus.

Despite changes in microglia and astrocyte populations, there was no evidence of amyloid plaque deposition in male or female LOAD2 or LOAD2.*Mthfr^677C>T^* mice (FIGURE 2D).

### Mthfr^677C>T^ modulates amyloid species and inflammatory cytokines in the brain

In the absence of plaque formation, levels of APP protein products were analyzed in brain lysates from 4-and 24-month-old cohorts. Both soluble and insoluble brain protein fractions were assessed using the Meso Scale Discovery (MSD) platform (see methods). As designed, only human Aβ species were detected in mice expressing humanized APP (FIGURE 3A). Levels of Aβ1-38 were below detection thresholds in all mice tested, in both soluble and insoluble protein lysate preparations. Both Aβ1-40 and Aβ1-42 levels increased with age in the insoluble fractions, with female mice consistently showing increased levels compared to males. Both young and aged female LOAD.*Mthfr^677C>T^*mice showed increased levels of combined Aβ1-40 and Aβ1-42 species compared to female LOAD2 controls, suggesting that the presence of *Mthfr^677C>T^*may impact APP processing or amyloid clearance in a sex-dependent manner. However, Aβ1-42/A1-β40 (42/40) ratios were not different between LOAD2 and *LOAD2.Mthfr^677C>T^* (SUPP FIGURE 3A) suggesting the increased levels of Aβ1-40 and Aβ1-42 may not predispose LOAD.*Mthfr^677C>T^* mice to cerebral amyloid angiopathy (CAA).

**FIGURE 3:**
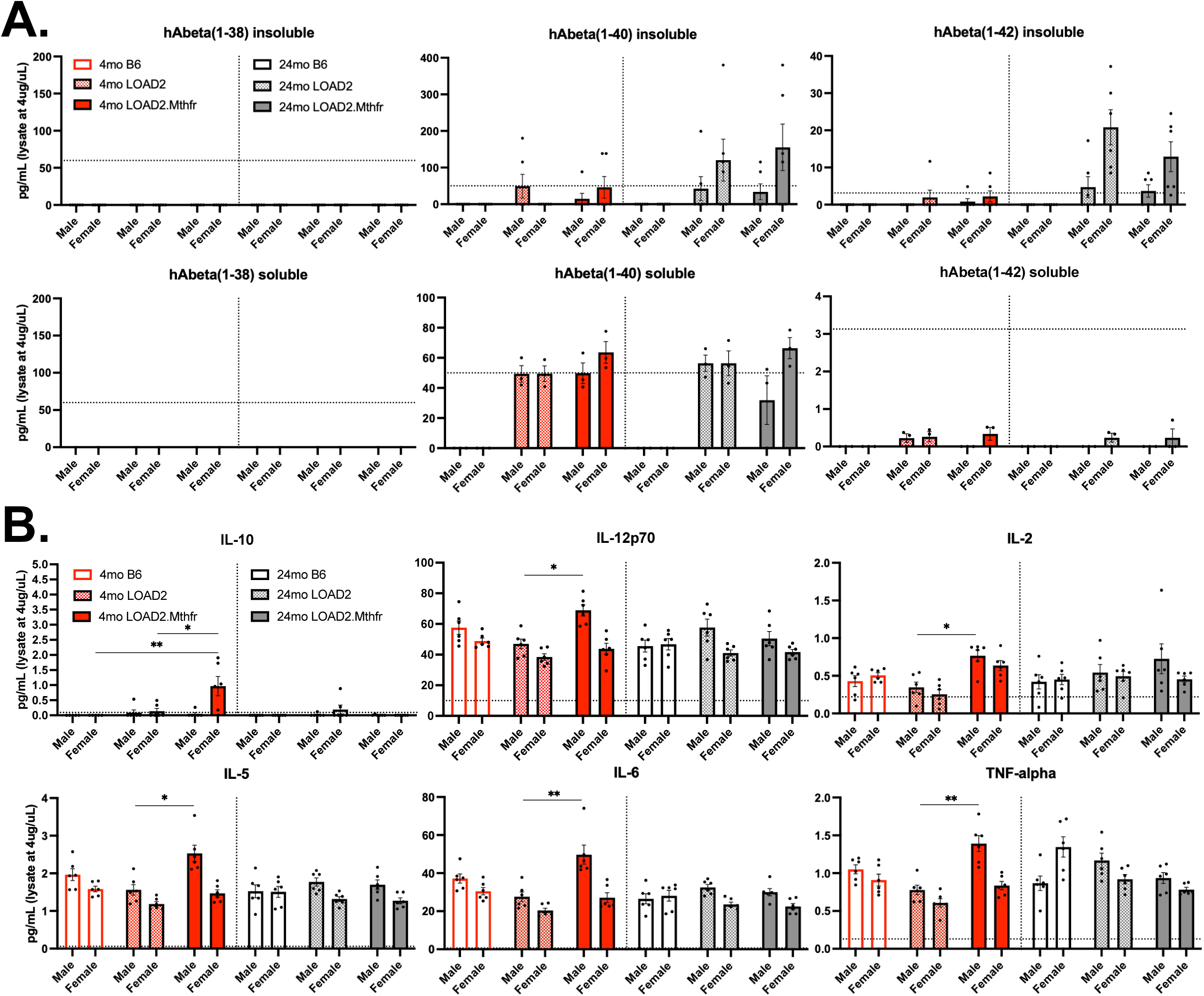
Inflammatory cytokine and human amyloid beta levels in brains of young and old LOAD animals. Young and aged C57BL/6J (B6), LOAD2, and LOAD2.*Mthfr^677C>T^*brain lysate analysis by MSD multi-plex assays. Concentrations of insoluble and soluble human amyloid species 1-38, 1-40, or 1-42 (A). Panel of interleukins (IL-10, −12p70, −2, −5, −6) and TNF-alpha depicting expression of soluble proteins in brain tissue (B). Sample size n≥5. Significant statistical difference determined within sex groupings by ANOVA: *p<0.05, **p<0.01, ***p<0.001.

Given the differences observed in microglia and astrocyte numbers, inflammatory cytokines and chemokines were assessed in brain lysates. Interferon (IFN)-gamma, IL-1beta, IL-4, and KC-GRO (CXCL1) expression levels in the brain were not significantly changed by age or genotype (SUPP FIGURE 3B). However, IL-10 levels were highest in young female LOAD2.*Mthfr^677C>T^*mice compared to young controls (FIGURE 3B). Similarly, IL-12p70, IL-2, IL-5, IL-6, and TNF-alpha levels were highest in young male LOAD2.*Mthfr^677C>T^*mice, greater than LOAD2 controls. We also observed, compared to B6, decreased soluble brain TNF-alpha in LOAD2 but not LOAD2.*Mthfr^677C>T^*mice. Collectively, these data suggest the Mthfr^677C>T^ variant modifies the inflammatory environment in the brain.

### Mthfr^677C>T^ has no significant effect on clinically relevant plasma biomarkers

Individual levels of Aβ species in blood plasma were not changed by age or genotype (FIGURE 4A; SUPP FIGURE 4A), but 24-month-old LOAD2.*Mthfr^677C>T^* male mice showed a reduced Aβ1-42/Aβ1-40 (42/40) ratio compared to the male LOAD2 group. As reported previously in LOAD2 [24], neurofilament-light-chain (Nf-L) in plasma increased with age (FIGURE 4B) irrespective of genotypes. Females generally showed higher levels of Nf-L, and male LOAD2.*Mthfr^677C>T^* mice showed the smallest increase with age. Inflammatory markers were also tested in the plasma, revealing decreased IL-4 levels in young female LOAD2.*Mthfr^677C>T^*mice compared to controls (FIGURE 4C). Age-related increases in IL-10 (SUPP FIGURE 4G) and TNF-alpha (SUPP FIGURE 4I) were noted across all genotypes, sexes, and treatments but did not reach significance. Confirming our previous data showing an increase in plasma homocysteine in *Mthfr^677C>T^* mice, LOAD2.*Mthfr^677C>T^*mice showed more plasma homocysteine than B6 and LOAD2 (FIGURE 4D)

**FIGURE 4:**
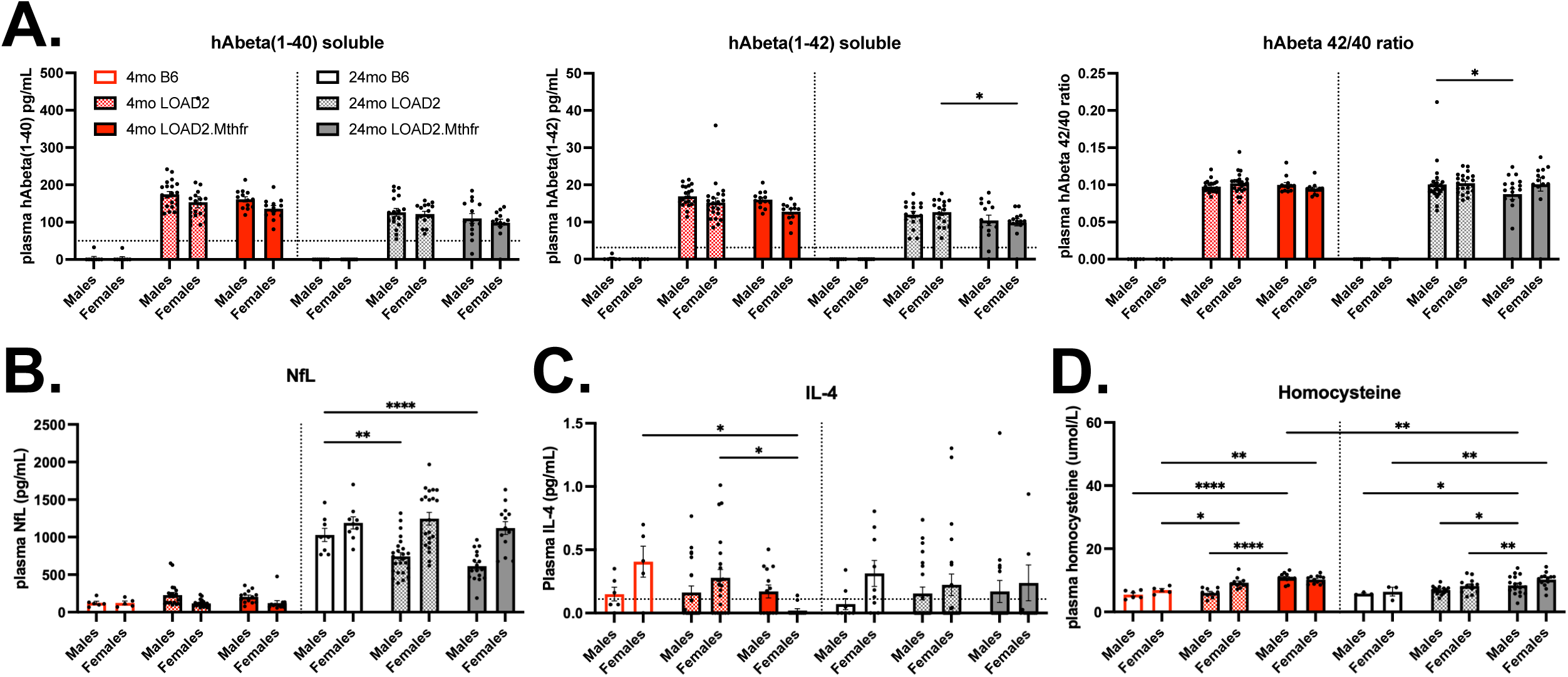
Soluble human amyloid beta, Nf-L, IL-4, and homocysteine levels in blood plasma of young and old LOAD mice. Young (4 month) and aged (24 month) C57BL/6J (B6), LOAD2, and LOAD2.*Mthfr^677C>T^* plasma analyzed by MSD multi-plex assays for levels of human amyloid beta species 1-40, 1-42, and 42/40 ratios (A). Plasma levels of neurofilament light-chain (Nf-L) (B), interleukin 4 (IL-4) (C), and homocysteine (D). Sample size n≥5. Significant statistical difference determined within sex groupings by ANOVA: *p<0.05, **p<0.01, ***p<0.001.

### Linear regression analyses reveal associations of phenotypes with genotype, sex, and age

To model the relationships between phenotypes (e.g. frailty, neuron counts, etc.,) and experimental variables (e.g. age, sex, and genotype) we performed linear regression analysis. Frailty index scores were positively associated with both age and *Mthfr^677C>T^* expression (FIGURE 5A), whereas no such relationships were identified for percentage correct in spontaneous alternation task in the Y maze (FIGURE 5B). As expected, linear regression analysis showed strong significance between levels of Aβ species (particularly Aβ1-40 and Aβ1-42) expression and genotype (LOAD2 and LOAD2.*Mthfr^677C>T^*) in both plasma (FIGURE 5C) and brain (FIGURE 5D). Interestingly, both LOAD2 and LOAD2.*Mthfr^677C>T^* were also associated with decreased IL-10, TNF-alpha, and interferon gamma in plasma (FIGURE 5C). Both LOAD2 and LOAD2.*Mthfr^677C>T^*were also significantly associated with plasma homocysteine levels, with LOAD2.*Mthfr^677C>T^* showing the strongest association (FIGURE 5C). Age at 24 months was a significant factor in plasma neurofilament and cytokine levels. In the brain, increased age was tied to decreased surface area and cell counts for both cortical and hippocampal neurons (NeuN+), astrocytes (GFAP+), and microglia (Iba1+) (FIGURE 5E).

**FIGURE 5:**
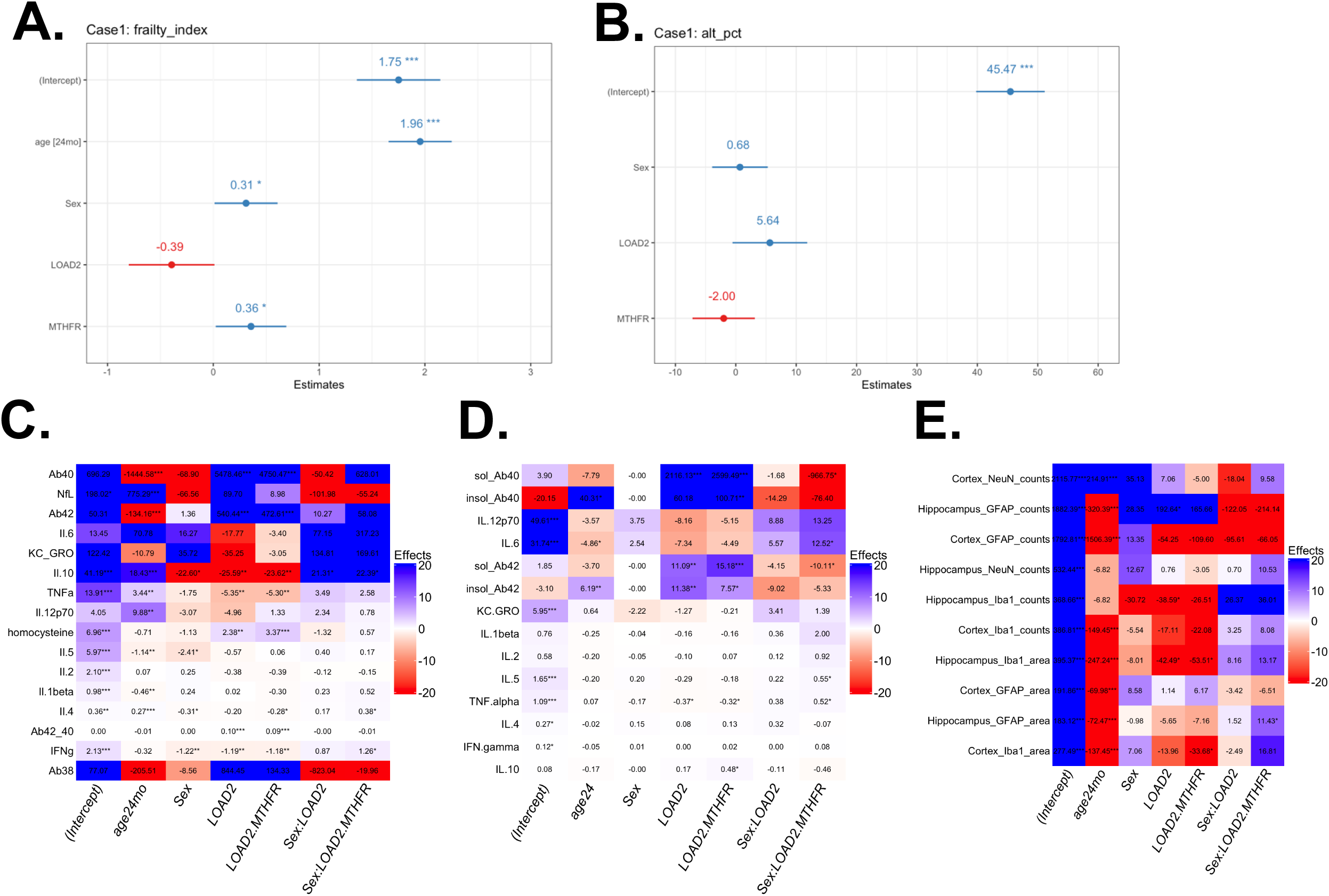
Individual risk factor analysis of phenotypes observed, and biomarker levels reveals isolated impact of AD risk factors on animal health. Cross sectional cohorts of aged (24-month) LOAD2 and LOAD2.*Mthfr^677C>T^*. Linear regression analysis of individual risk factors (age, sex, genetic background, *Mthfr*-expression) effect on cumulative frailty index biometric score (A), spontaneous alternation (Y-maze) behavioral assay performance (B), blood-based biomarker protein levels (C), brain-based biomarkers (D), and immunohistochemical cellular biomarkers (E). Sample size n≥5. Significant statistical difference determined within sex groupings by linear regression: *p<0.05, **p<0.01, ***p<0.001.

### Brain transcriptomics suggest LOAD2.Mthfr^677C>T^ present an accelerated aging phenotype

To further examine the value of LOAD2.*Mthfr^677C>T^*as a preclinical model for LOAD, we performed transcriptional and proteomic profiling on brain samples from 4-, 12-, 18-and 24 months old male and female LOAD2 and LOAD2.*Mthfr^677C>T^* mice, and 4-and 24 months old male and female B6 mice. To identify upregulated and downregulated differentially expressed genes (DEGs), data from LOAD2.*Mthfr^677C>T^*was compared to either B6 mice (effect of LOAD2 alleles and *Mthfr^677C>T^*variant) or LOAD2 mice (effect of *Mthfr^677C>T^* variant) (FIGURE 6A). Both comparisons revealed sex-and age-dependent differences. Interestingly, the number of DEGs comparing LOAD2.*Mthfr^677C>T^* to B6 was greater at 4 months of age than 24 months of age (FIGURE 6A). This suggests LOAD2.*Mthfr^677C>T^*may display an accelerated or non-linear aging signature. Among the downregulated DEGs at 24 months were genes associated with activity in AD including *Trem2, Itgax,* and *Apoe*, particularly in females (FIGURE 6B). These findings may be due to the presence of *Trem2^R47H^*allele in LOAD2.*Mthfr^677C>T^* mice that is known to suppress microglia function. This finding also correlates with the reduced numbers of microglia we observed by immunofluorescence (FIGURE 2). Comparing LOAD2.*Mthfr^677C>T^*to LOAD2 mice, DEGs were only identified in 4-or 18-month samples, suggesting the effect of the *Mthfr^677C>T^*on the brain throughout aging is not linear (FIGURE 6A).

**FIGURE 6:**
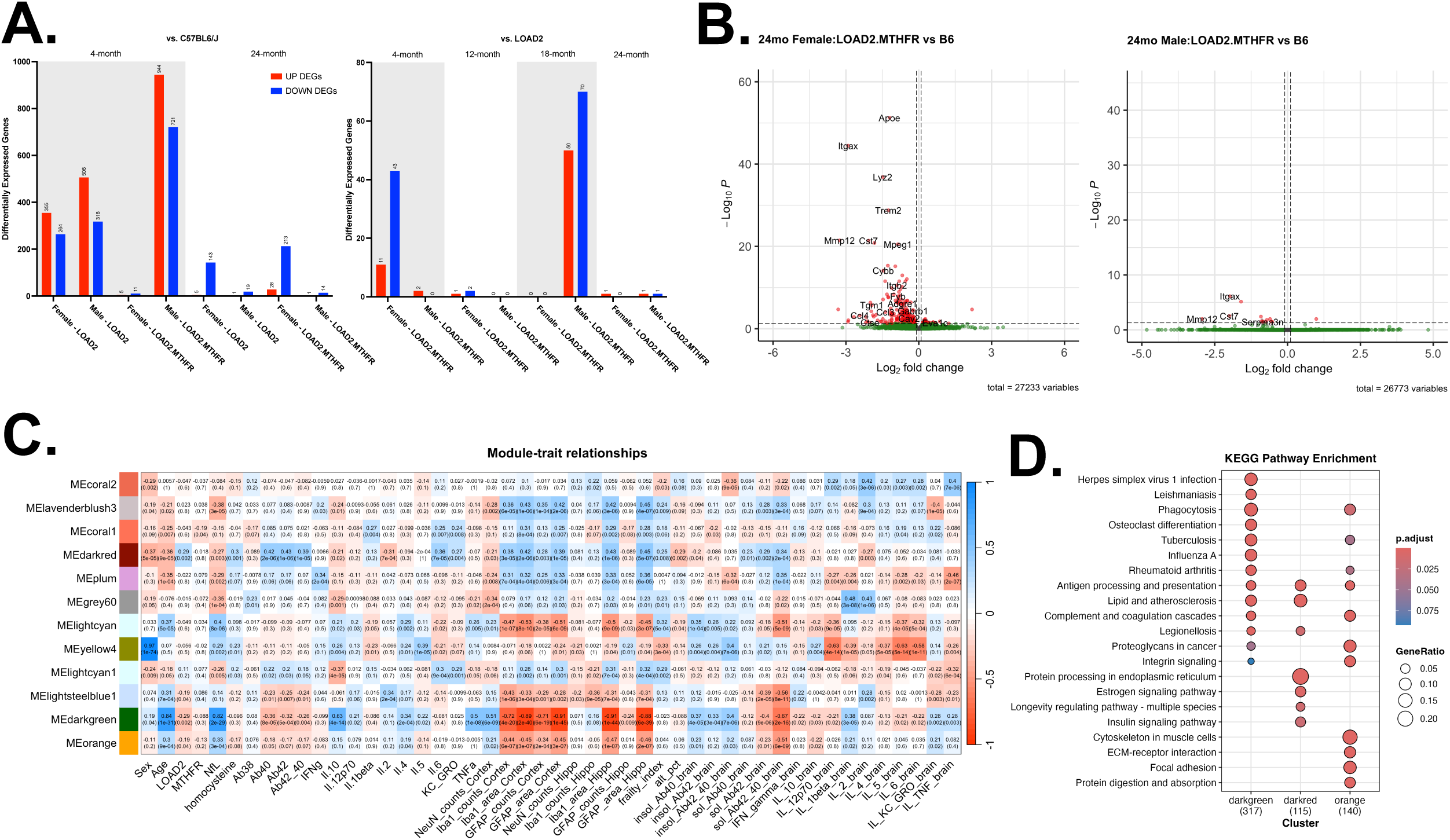
Module-trait relationships identify potential biomarkers changes associated with expression. Plots of differentially expressed genes (DEGs) in LOAD2 and LOAD2.*Mthfr^677C>T^* compared to C57BL/6J (B6) and LOAD2 from 4 months of age to 24 months (A). Volcano plots of DEGs using DESeq2 (B). Differences in gene transcription within eigengene modules were compared to AD risk factors, phenotypes, and biomarkers in the brain. Matrix showing relationship between modules and risk factors (sex, age, LOAD2 genetic background, *Mthfr^677C>T^* expression), inflammatory cytokine levels in blood or brain, cellular profiles (neurons, microglia, astrocytes) in the hippocampus or cortex, and levels of soluble or insoluble hAbeta species (1-38, 1-40, 1-42) (C). Enriched processes for MEDarkgreen, MEDarkRed, and MEOrange (D). Data from cross sectional cohorts of aged (24-month) LOAD2 and LOAD2.Mthfr 677C>T. Sample size n≥5. Significant statistical difference determined within sex groupings by linear regression: *p<0.05, **p<0.01, ***p<0.001.

Weighted gene co-expression network analysis (WGCNA) [52] identified relationships between gene expression modules and phenotypes (FIGURE 6C). No module was strongly correlated with the *Mthfr^677C>T^*variant but two modules (MEdarkgreen, MEorange) were positively correlated with age, and Nf-L, but negatively correlated with plasma Aβ species and cell counts in the brain. Interestingly, the correlations were inverted for MEdarkred. Genes in MEdarkgreen are enriched for immune-related processes, MEorange for immune-and vascular-related processes, and MEdarkred for aging-, cell stress-, and insulin signaling-related processes (FIGURE 6D) suggesting these processes may be impaired in LOAD2*.Mthfr^677C>T^* mice.

To assess similarity of transcriptomic signatures in *LOAD2.Mthfr^677C>T^* mice to human AD we aligned all gene expression changes (logFC values) from the 24-month *LOAD2.Mthfr^677C>T^* to B6 comparison with AMP-AD gene modules that represent molecular changes in human AD (SUPP FIGURE 5A) [55]. We also included transcriptomic data from 18 months old LOAD2 mice fed a high fat diet that we had previously characterized as a useful preclinical model [35]. Organization of gene expression signatures into AMP-AD expression modules [55] highlight age, sex, and genotype-driven alterations within consensus clusters based on cellular systems in the brain. Differential gene expression data from experimental mouse cohorts are contrasted against disease-driven gene expression changes from human datasets. As expected due to the lack of amyloid pathology that is known to increase expression of immune genes, both LOAD2 HFD and LOAD2.*Mthfr^677C>T^*mice showed negative correlations with modules in Consensus Cluster B (enriched for plaque-related immune system genes). However, both models showed positive correlations with modules in Consensus Cluster C (Neuronal System) but in a sex-dependent fashion (SUPP FIGURE 5A). Specifically, LOAD2 HFD males were more strongly correlated than LOAD2 HFD females. Conversely, LOAD2.*Mthfr^677C>T^*females were more strongly correlated with neuronal modules than their male counterparts. LOAD2.*Mthfr^677C>T^* females also showed significant positive correlations with Consensus Cluster E (Organelle Biogenesis, Cellular stress response) (SUPP FIGURE 5A). The negative correlations with immune system-related modules, but positive correlations with non-immune related modules was further evident when comparing gene expression changes (logFC values) to AD subtypes that can be categorized as inflammatory or non-inflammatory (SUPP FIGURE 5B-C). When considering B6 as the control, positive correlations were primarily observed between LOAD2 or LOAD2.*Mthfr^677C>T^* and non-inflammatory subtypes (SUPP FIGURE 5B). However, when considering LOAD2*.Mthfr^677C>T^*compared to LOAD2, the *Mthfr^677C>T^* variant induces an age-dependent switch from a non-inflammatory to an inflammatory phenotype that is accelerated in females (SUPP FIGURE 5C).

To further evaluate relevance of the LOAD2.*Mthfr^677C>T^*model to human AD and to identify signatures specific to the *Mthfr^677>T^* variant, we aligned gene expression changes (logFC values) from LOAD2.*Mthfr^677C>T^* compared to LOAD comparisons from age groups (4-, 12-, 18-and 24 months) (FIGURE 7). Strong gene expression correlations emerged at 24 months of age in non-sex-stratified comparisons between LOAD2 and LOAD2.*Mthfr^677C>T^* mice (FIGURE 7A). When stratified by sex, female LOAD2.*Mthfr^677C>T^* mice compared to human AD modules are strongly correlated at 12 months with the ‘Cell Cycle, Consensus Cluster D’, and strongly correlated at 18-and 24 months with the ‘Neuronal, Consensus Cluster C’ and ‘Organelle Biogenesis, Consensus Cluster E’ (FIGURE 7B). Male LOAD2.*Mthfr^677C>T^* mice compared to the human AD modules primarily show correlations at 12 months within the ‘Neuronal, Consensus Cluster C’ and ‘Organelle Biogenesis, Consensus Cluster E’ (FIGURE 7C). Finally, non-sex stratified comparisons with biodomains developed as part of the TREAT-AD program, revealed age-dependent correlations in gene expression signatures across all categories, including lipid metabolism, myelination, synapse, and vasculature (FIGURE 7D).

**FIGURE 7:**
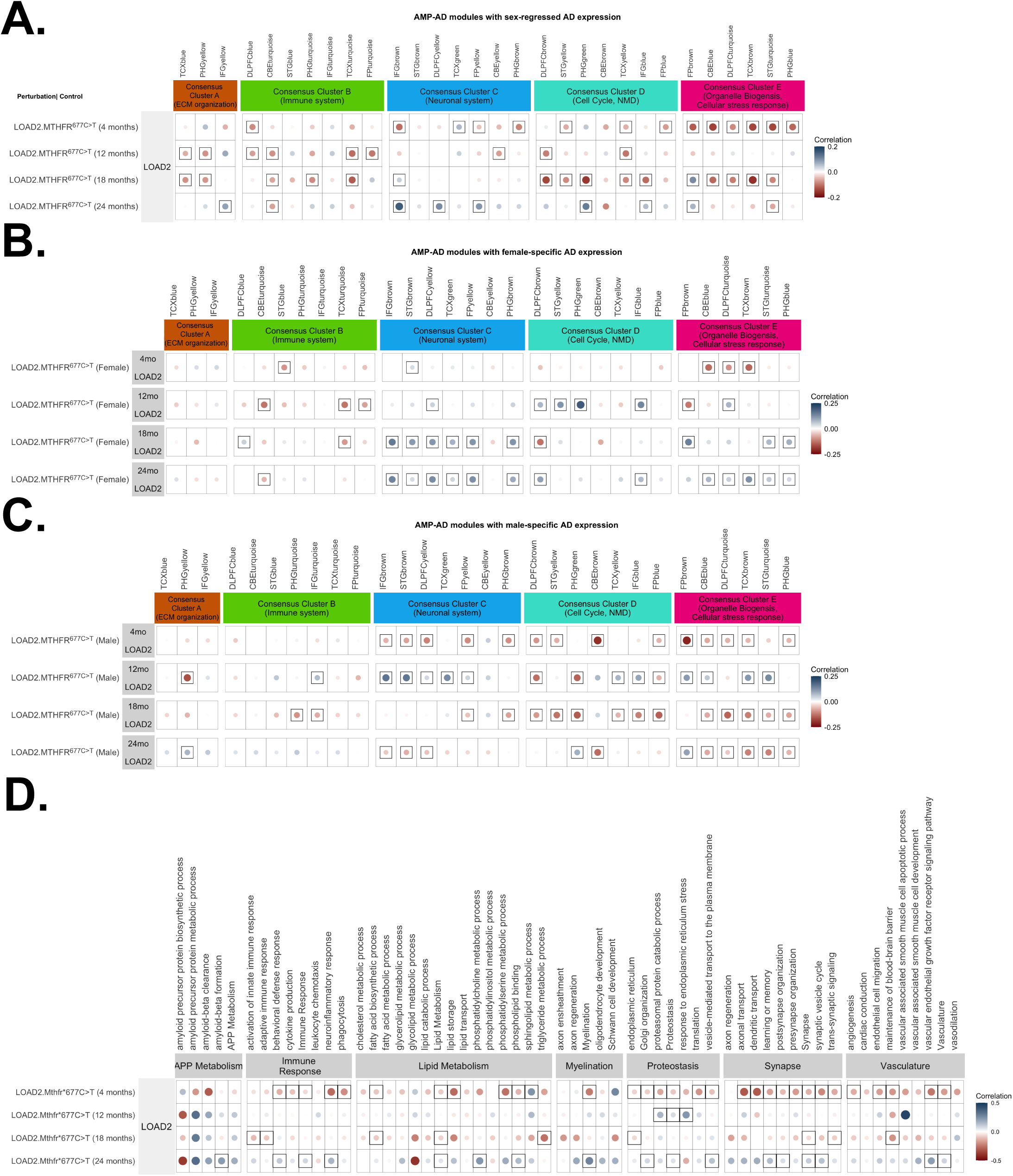
Transcriptomic profiling of brain tissue from LOAD animals with and without expression of *Mthfr* risk allele. Gene expression driven by *Mthfr^677C>T^* organized into AMP-AD expression modules, in both males and females (A), compared to human LOAD patient expression. Sex-stratified gene expression organized into AMP-AD modules and correlated to female-(B) or male-specific (C) AD vs. control human patient expression. Gene expression organized into TREAT-AD biodomain representation for both sexes (D). Nineteen distinct biological domains have been identified and are defined using sets of GO terms that align with the endophenotype. Cross sectional cohorts of C57BL/6J (B6), LOAD2 and LOAD2.*Mthfr^677C>T^*. Sample size n≥5. Circles denote positive (blue) or negative (red) correlations; color intensity and circle size reflect the correlation magnitude, with significant correlations (p < 0.05) indicated by frames.

### Mthfr^677C>T^ predicted to mediate cerebrovascular and white matter health

Previous work has suggested that protein analysis may better represent changes in the brain related to human AD than transcriptomics [50]. Therefore, to test this, and to further evaluate LOAD2.*Mthfr^677C>T^* as a preclinical model, we performed proteomics on the same brains from LOAD2.*Mthfr^677C>T^* mice and controls that we performed transcriptomic analyses on. Differentially expressed proteins (DEPs) were calculated comparing LOAD2.*Mthfr^677C>T^*to either B6 (at 4-and 24 months of age; to determine the value of the model for preclinical studies) or LOAD2 (at 4-, 12-, 18-, and 24 months of age; to identify *Mthfr^677>T^* variant effects). Results showed that there were indeed more DEPs (FIGURE 8A) than DEGs (FIGURE 6A). At 4 months, there were more DEPs comparing LOAD2.*Mthfr^677C>T^* to LOAD than B6. Similar numbers of DEPs were observed for both comparisons at 24 months of age. Considering the LOAD2.*Mthfr^677C>T^* to LOAD2 comparison, the number of DEPs was similar between 4-and 12 months, with a significant increase in upregulated DEPs at 18 months. There were fewer DEPs at 24 months of age, further supporting some aspects of aging may be accelerated or non-linear in the presence of the *Mthfr^677C>T^*variant. DEPs also varied between sexes suggesting sex-specific actions of the *Mthfr^677C>T^* that we have previously reported [33]. DEPs we compared to the proteins expressed in human AD compared to controls (Emory DEPs) [50]. Numbers of Emory DEPs were relatively consistent across all ages and comparisons (FIGURE 8A). One key protein that was differentially expressed in male, but not female, LOAD2.*Mthfr^677C>T^* compared to B6 at 24 months was SMOC1 (FIGURE 8B-C) that has been associated with cerebrovascular dysfunction [61], a result that fits with the *Mthfr^677C>T^* inducing vascular-related deficits.

**FIGURE 8:**
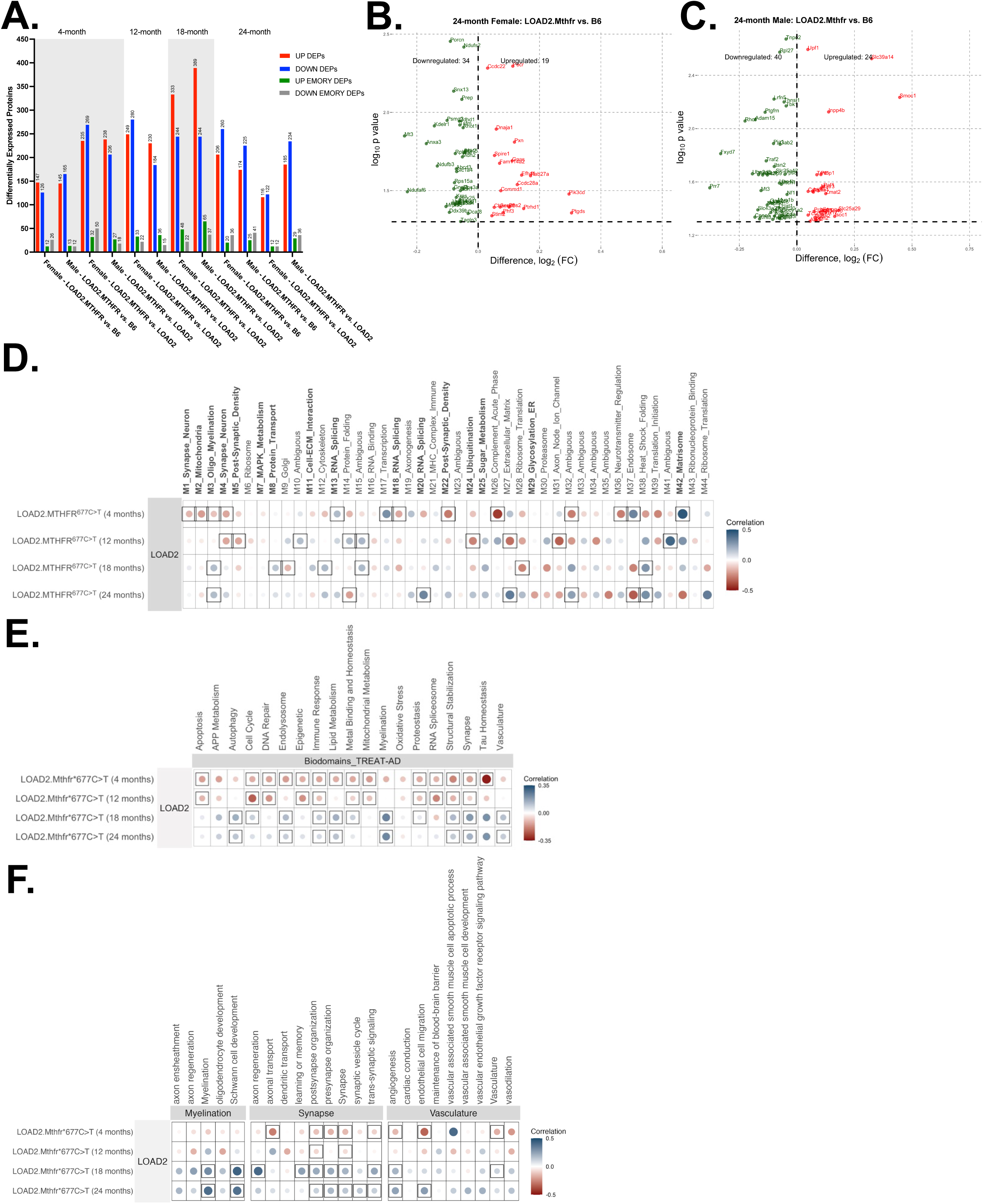
*Mthfr^677C>T^*expression alters proteomic signature of aged LOAD mice brains towards expression observed in human LOAD. Cross sectional cohorts of untreated C57BL/6J, LOAD2 and LOAD2.*Mthfr^677C>T^*. Table of differentially expressed proteins (padj < 0.05) (A). Volcano plots of DEPs classified by EMORY module designation in 24-month-old LOAD2.*Mthfr^677C>T^* males and females vs. B6 controls (B-C). Alignment of mouse proteomic data to Emory human modules showing Pearson correlation coefficients between protein expression change in in LOAD2.*Mthfr^677C>T^* mice relative to LOAD2 at different ages and protein expression changes in human AD subjects (log fold change for cases versus controls) (D). Blue and red circles represent positive and negative correlations, respectively; circle size and color intensity reflect the correlation magnitude. Significant associations (p<0.05) are highlighted with black frames. Alignment of mouse proteomic data to TREAT-AD Biodomains (E) and relevant subdomains (F). Circles represent positive (blue) or negative (red) correlations; circle size and color intensity reflect correlation strength, with significant correlations (p < 0.05) framed.

To dissect the contribution of the *Mthfr^677C>T^* variant to AD, protein expression changes in LOAD2.*Mthfr^677C>T^*compared to LOAD at 4-, 12-, 18-and 24 months of age, were aligned to the disease associated human LOAD modules from the Emory study [50] (FIGURE 8D). Positive correlations were present at all ages, with fewer negative correlations with age. Among the positively correlated modules at 4 months was the M42_matrisome module that has been associated with cerebrovascular changes [50]. Among the positively correlated modules at 24 months of age was the M3_Oligo_Myelination module that is associated with white matter health. These correlations suggest the *Mthfr^677C>T^* variant is a useful tool to study vascular contributions to cognitive impairment and dementia (VCID). To assess this, protein expression changes in LOAD2.*Mthfr^677C>T^*compared to LOAD at 4-, 12-, 18-and 24 months of age were aligned to the biodomains developed by the TREAT-AD consortium [60] (FIGURE 8E-F). The greatest number of positive correlations was observed at 18 and 24 months of age, while there were only negative correlations in the 4-and 12-month-old comparisons. This shows the age-dependency of the *Mthfr^677C>T^* variant. Myelination, Synapse and Vasculature biodomains were strongly correlated at 18-and 24 months of age (FIGURE 8E). Subdomain analysis suggests perturbations to synapses rather than axons, and angiogenesis and endothelial cell migration (FIGURE 8F), further supporting the use of the *Mthfr^677C>T^* variant to study VCID.

## DISCUSSION

Data presented here introduces the LOAD2.*Mthfr^677C>T^*mouse model and provides evidence that the presence of the *Mthfr^677C>T^*variant increases human AD relevance. *In vivo,* longitudinal behavioral and biometric analysis revealed no significant phenotypes due to the LOAD2 alleles or LOAD2 in concert with *Mthfr*^677C>T^. Neuropathology reveals that neither LOAD2 nor LOAD2.*Mthfr^677C>T^*mice develop ThioS-positive amyloid plaques, which may explain the absence of cognitive phenotypes. However, analyses of brain lysate showed elevated soluble and insoluble amyloid species in both LOAD2 and LOAD2.*Mthfr^677C>T^*, most prominently in females; this demonstrates the effect of hAbeta in the model and suggests a sex-dependent clearance or production mechanism. Further immunostaining demonstrated LOAD2-dependent changes within glial cell populations and analysis of brain lysate and plasma reveal *Mthfr^677C>T^*-dependent immune dysregulation. Transcriptomic and proteomic analysis of mice at multiple timepoints demonstrate sex-age-and genotype-dependent expression changes, including module-trait relationships. At 18-and 24 months of age, transcriptomic and proteomic data suggest LOAD2 and LOAD2.*Mthfr^677C>T^*show expression changes similar to those seen in human AD, with the LOAD2.*Mthfr^677C>T^*mice showing the strongest correlations. These data suggest that the *Mthfr^677C>T^* variant may amplify relevant AD phenotypes, possibly by modulating early cerebrovascular and immune responses.

Methylenetetrahydrofolate reductase (MTHFR) is a key rate-limiting enzyme in the folate and methionine cycles. The amino acid homocysteine is an important substrate produced as part of the metabolism of methionine. However, excess production can result in hyperhomocysteinemia, which is linked to cardio-and cerebrovascular dysfunction [32]. The *MTHFR^677C>T^* polymorphism codes for a heat-sensitive variant characterized by reduced MTHFR enzymatic activity resulting in increased plasma homocysteine levels [62–64]. At a cellular level, elevated plasma homocysteine promotes vascular inflammation and endothelial dysfunction [65–67] and is associated with poor cognitive function [68], Alzheimer’s disease [69], and vascular dementia [70–72]. Reduced MTHFR activity can also reduce folate and B12 production, as well as alter DNA methylation capabilities, all of which are important to maintain cognitive health [73–75]. While the presence of the *Mthfr^677C>T^* variant does not directly cause Alzheimer’s disease and related dementias, its relationship to additive systemic damage is relevant as up to 40 percent of people can be heterozygous and up to 20 percent can be homozygous for the T risk allele [32, 76].

The primary goal of the IU/JAX/PITT MODEL AD Center is to develop mouse models that more accurately represent the complexity of LOAD in a diverse population. Widespread evidence in the field suggests that vascular dysfunction is a critical factor for disease development and progression [77–79], but few mouse models have been created to specifically study genetic contributions to cerebrovascular health in the context of AD. Members of our group previously generated a mouse to model the *Mthfr^677C>T^* risk variant in isolation. Results showed elevated plasma homocysteine and reduced MTHFR enzymatic activity in liver and brain, as well as vascular deficits in brain [33] and retina [80], suggesting its clinical translatability. To further examine the role of *Mthfr^677C>T^*on a background of AD risk, the LOAD2*.Mthfr^677C>T^* model was created by the IU/JAX/PITT MODEL-AD Center by incorporating the *Mthfr^677C>T^*allele with the LOAD2 alleles (*hAbeta/APOE4/Trem2*R47H*).

All aged mice predictably scored higher on the frailty assay, with 18-month-old female LOAD2.*Mthfr^677C>T^* demonstrating the highest scores (FIGURE 1). While these data are not statistically significant, they suggest a sex-dependent baseline susceptibility that we observe throughout this study. Male LOAD2.*Mthfr^677C>T^* mice at 24 months of age did show increased rearing and decreased sum margin time in the open field assay, suggesting possible hyperexcitability (FIGURE 1). All 24-month-old male mice showed an increase in NeuN positive cells in the subiculum compared to females (FIGURE 2), which could contribute to a hyperexcitable phenotype.

We did not observe neurodegeneration or plaque deposition in the mice at any age (FIGURE 2; SUPP FIGURE 1; SUPP FIGURE 2). Although examining the effects of humanized Abeta is valuable in order to increase human-relevant AD pathology, adding *Mthfr^677C>T^* to models of known amyloid or tau drivers (e.g. *hAbeta^SAA^* or *MAPT*) will likely drive more visible pathological phenotypes. Similarly, in mice carrying the *Mthfr^677C>T^*variant alone, increased numbers of IBA1-expressing microglia and GFAP-expressing astrocytes were observed in several cortical regions, almost exclusively in males [33]. Initially numbers of GFAP-expressing astrocytes were decreased in 4-month-old male LOAD2.*Mthfr^677C>T^*mice (SUPP FIGURE 2), but by 24 months, the numbers had normalized. Alternatively, aged female LOAD2 and LOAD2.*Mthfr^677C>T^*mice showed an increase in number of astrocytes (FIGURE 2) and a decrease in cell area (SUPP FIGURE 1), suggesting the cells are in an activated state. However, microglial cell density was reduced in both LOAD2 and LOAD2.*Mthfr^677C>T^* mice at young and aged timepoints with a stronger reduction in females (FIGURE 2; SUPP FIGURE 2).

While we did not observe dramatic phenotypes in the LOAD2*.Mthfr^677C>T^*mice, an interesting trend is emerging. In the present study, 4-month-old LOAD2*.Mthfr^677C>T^* mice showed a decrease in cortical microglial counts (SUPP FIGURE 2), as well as significant changes in cytokine/chemokine expression regulation that normalize with age. Female LOAD2*.Mthfr^677C>T^* mice at 4 months showed significantly higher expression of brain IL-10, an anti-inflammatory cytokine that is upregulated in response to CNS injury or neuroinflammation (FIGURE 3). Conversely, the same group of mice showed a significant decrease in expression of plasma IL-4, (SUPP FIGURE 4) which can represent an imbalance in immune function and anti-inflammatory regulation. In our previous studies, *Mthfr^677C>T^* alone demonstrated aberrant vascular or immune signatures by 6 months of age, some of which appeared in control mice at later ages [33]. This suggests that vascular and immune dysregulation may be influenced by a decrease in MTHFR enzyme activity at an early age, priming the microenvironment to respond poorly to additional cellular stressors. Chronic inflammation can result in an early aging, or inflammaging, phenotype, with cells demonstrating an increased likelihood of entering a senescent stage [81].

By examining the gene and protein expression correlations between the LOAD mice and human AD patients, we observed that aged LOAD2*.Mthfr^677C>T^* show more correlations in almost every biological category, including myelination, immune response, cellular stress, and vasculature, suggesting *Mthfr^677C>T^*is driving the model towards a human AD ‘state’. Because we noted cerebrovascular-relevant signatures in the AMP-AD disease-associated modules and the TREAT-AD biodomains, more precise vascular phenotyping is warranted in the future.

## CONCLUSION

MODEL-AD continues to create strains that develop endophenotypes of LOAD for preclinical testing. While they represent more subtle dysfunction, changes observed due to *Mthfr^677C>T^* may provide a promising way to examine additive risk, and developing additional polygenic models is in progress. Ultimately, this work will aid in the identification of novel therapeutic approaches with the goal of reducing inflammatory and cerebrovascular-related pathology in ADRDs.

## Supporting information

Supplemental Figures images

## LIST OF ABBREVIATIONS

ADRD: Alzheimer’s disease and related dementias
LOAD: Late onset Alzheimer’s disease
MODEL-AD: Model Organism Development and Evaluation for Late-onset Alzheimer’s Disease
MTHFR: Methylenetetrahydrofolate reductase
TREAT-AD: Target Enablement to Accelerate Therapy Development for Alzheimer’s Disease
VCID: Vascular contributions to cognitive impairment and dementia

## DECLARATIONS

### Ethics approval and consent to participate

This study was approved by The Jackson Laboratory Institutional Animal Care and Use Committee (IUACUC).

### Consent for publication

Not applicable

### Availability of data and materials

Data is being made available through Synapse.

### Competing interests

The authors declare that they have no competing interests.

## Funding

This work was funded by the IU/JAX/PITT MODEL-AD Consortium (U54 AG054345). GH was supported by the Diana Davis Spencer Endowed Chair Research and GC was supported by the Bernard and Lusia Milch Endowed Chair.

## Authors’ contributions

Project design was performed by K.P.K., A.M.R., M.S., G.W.C., G.R.H. Experimentation was performed by K.P.K., Z.S., R.O., S.H., A.D., N.T.S. Data analysis/presentation was performed by K.P.K., R.S.P. Manuscript preparation was performed by K.P.K., A.M.R., R.S.P., G.W.C., G.R.H. All authors approved the submission.

## Acknowledgements

We gratefully acknowledge the contributions of the Genome Technologies, Center for Biometric Analysis, Neurobehavioral Phenotyping, Research Animal Facilities, Genetic Engineering Technologies, and Clinical Chemistry Staff and Services at The Jackson Laboratory in Bar Harbor, Maine for their expert assistance with the work described in this publication. The authors would like to acknowledge Martha Abbot for her outstanding efforts and support as the MODEL-AD administrator at The Jackson Laboratory during all phases of this project.

## Notes

### Competing Interest Statement

The authors have declared no competing interest.

