## Supplemental Figures text for "MTHFR*677C>T produces distinct prodromal disease signatures in a mouse model of late-onset Alzheimer’s disease"

**SUPPLEMENTAL FIGURE LEGENDS**

**SUPP FIGURE 1: Neuropathological assessment of aged brain tissue.** C57BL/6J (B6), LOAD2, and LOAD2.*Mthfr^677C>T^* 24-month-old cross-sectional immunohistochemistry. Representative 20X magnification images analyzed in each brain, cortex and hippocampus (A). DAPI+ counts (B). Average cell area of cells counted stained positive for Iba1 (microglia) (C) and GFAP (astrocytes) (D). Three separate sections per brain (bregma markers approximately -2.00 mm). Sample size n≥15. Significant statistical difference determined within sex groupings by ANOVA: *p<0.05, **p<0.01, ***p<0.001.

**SUPP FIGURE 2: Neuropathological assessment of young brain tissue.** 4-month-old C57BL/6J (B6), LOAD2, and LOAD2.*Mthfr^677C>T^* cross-sectional cohort immunohistochemistry of cortex and hippocampus. Total DAPI+ counts (A), GFAP+ (astrocytes) (B), and Iba1+ (microglia) (C). Representative 20X magnification images analyzed in each brain, cortex and hippocampus. Bregma markers approximately -2.00 mm. Sample size n≥15. Significant statistical difference determined within sex groupings by ANOVA: *p<0.05, **p<0.01, ***p<0.001.

**SUPP FIGURE 3: Interferon-gamma, interleukin-1-beta, interleukin-4, KC-GRO, and soluble human Abeta levels in young and old LOAD brain tissue.** Young and aged C57BL/6J (B6), LOAD2, and LOAD2.*Mthfr^677C>T^* brain lysate analysis by multi-plex MSD. Human amyloid beta 42/40 ratios (A) and mouse inflammatory (IFN-gamma, IL-1beta, IL-4, and KC-GRO) (B) measurements in soluble fractions of brain homogenate. Sample size n≥5. Significant statistical difference determined within sex groupings by ANOVA: *p<0.05, **p<0.01, ***p<0.001.

**SUPP FIGURE 4: Soluble human amyloid beta 1-38, inflammatory modulating cytokines, KC-GRO, and TNF-alpha levels in blood plasma of young and old LOAD mice.**

Young and aged C57BL/6J (B6), LOAD2, and LOAD2.*Mthfr^677C>T^* plasma analyzed by MSD multi-plex assays for levels of human amyloid beta species 1-38 (A). Plasma levels of interleukins 12p70 (IL-12p70) (B), IL-1beta (C), IL-2 (D), IL-5 (E), IL-6 (F), IL-10 (G), KC-GRO (H), and TNF-alpha (D). Sample size n≥5. Significant statistical difference determined within sex groupings by ANOVA: *p<0.05, **p<0.01, ***p<0.001.

**SUPP FIGURE 5: Gene expression correlations between LOAD animals and human patients.** Sex-stratified AMP-AD into 30 co-expression modules related LOAD pathology from human cohort study comparing effects of high-fat-diet (HFD), *Mthfr^677C>T^*, and/or age compared to C57BL/6J (B6) (A). These consensus clusters consist of a subset of modules which are associated with similar AD related changes across the multiple studies and brain regions. AD subtypes in ROSMAP, Mayo and MSBB cohort expression modules comparing LOAD2.*Mthfr^677C>T^* vs B6 (B) and LOAD2 (C) genotype controls at multiple ages. Inflammatory (ROSMAP_subtypeA, Mayo_subtypeA & B, MSBB_subtypeA) and non-inflammatory subtypes. Sample size n≥5. Circles denote positive (blue) or negative (red) correlations; color intensity and circle size reflect the correlation magnitude, with significant correlations (p < 0.05) indicated by frames.
