## Supplementary figures and images for "MTHFR*677C>T produces distinct prodromal disease signatures in a mouse model of late-onset Alzheimer’s disease"

### Supplemental Figures images

#### SUPP FIGURE 1

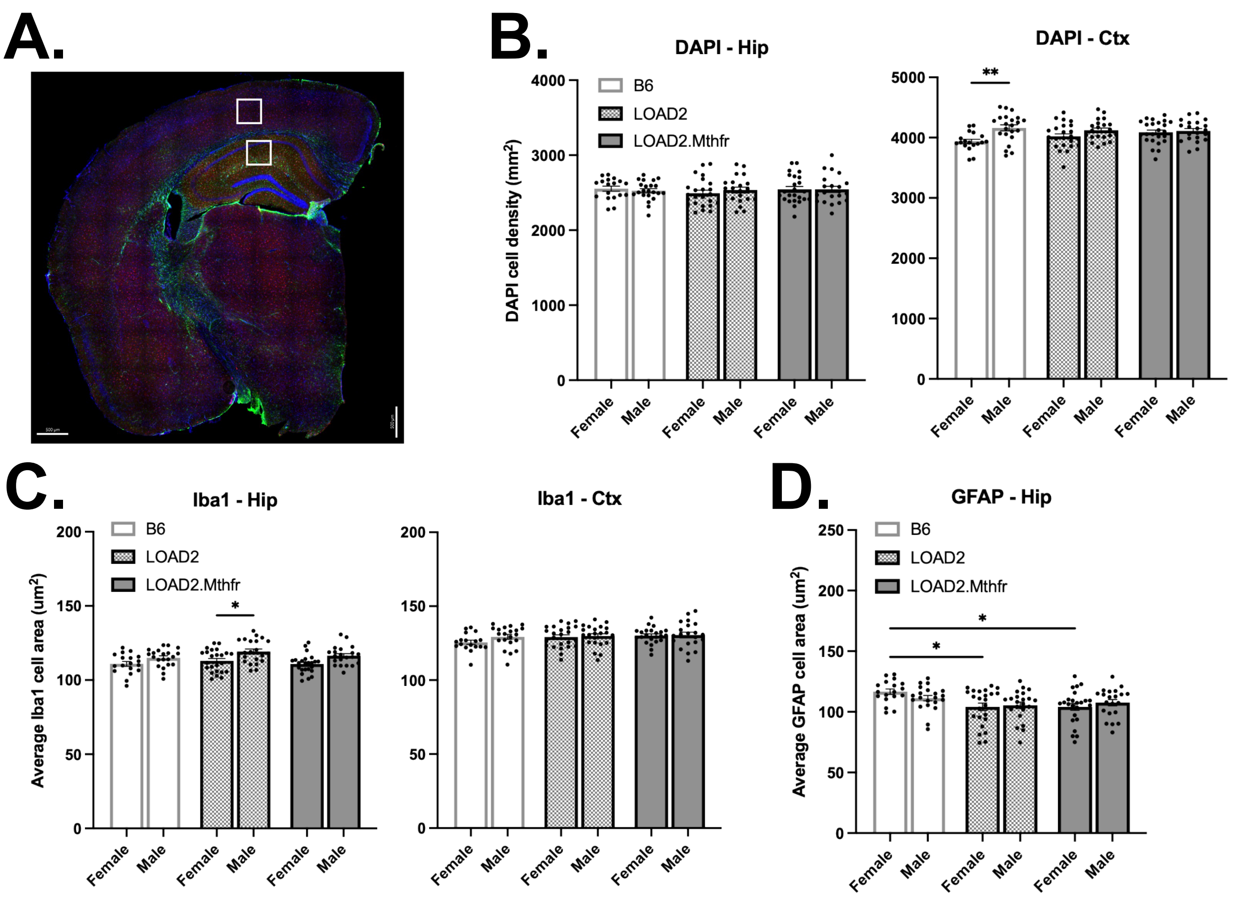

#### SUPP FIGURE 2

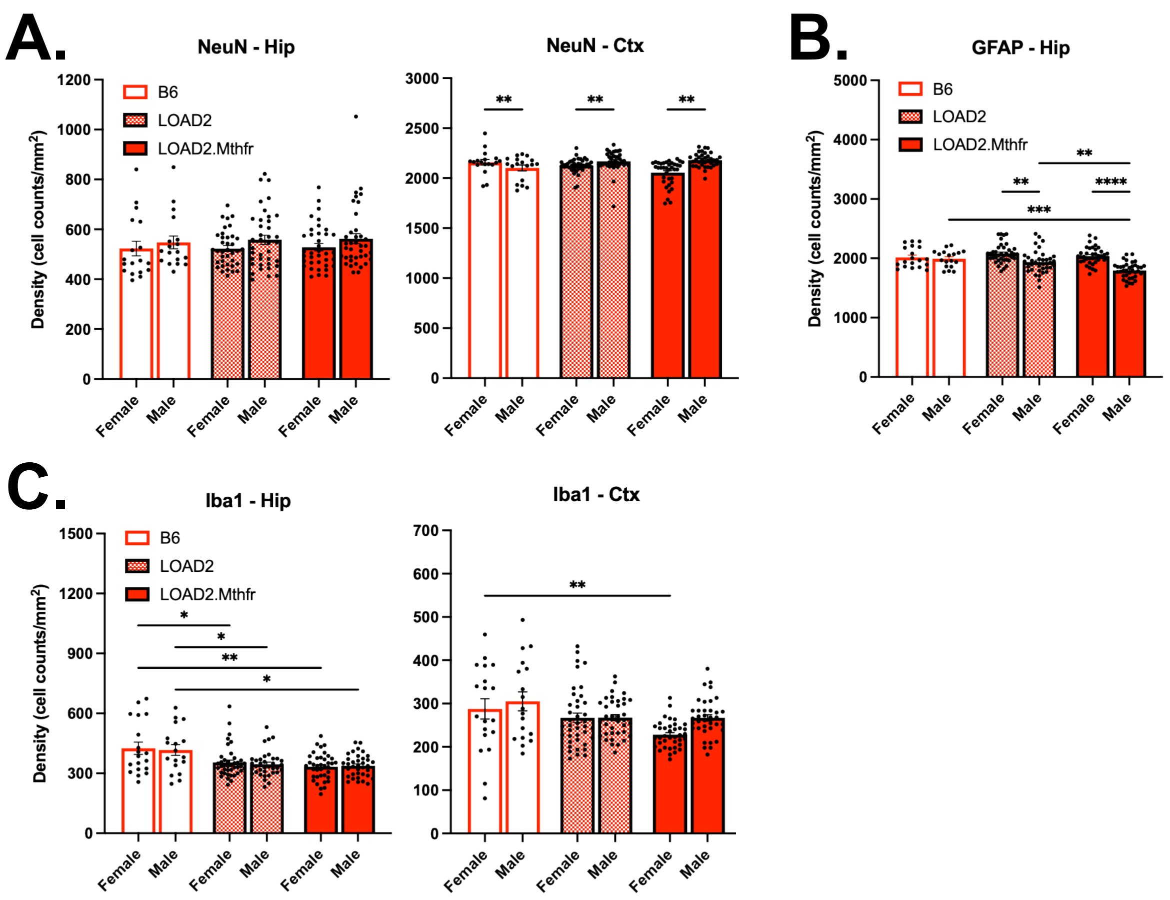

#### SUPP FIGURE 3

**A.**

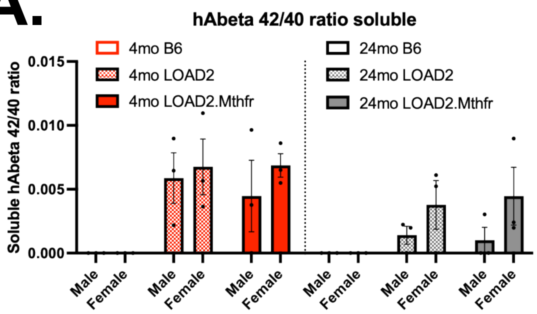


**B.**

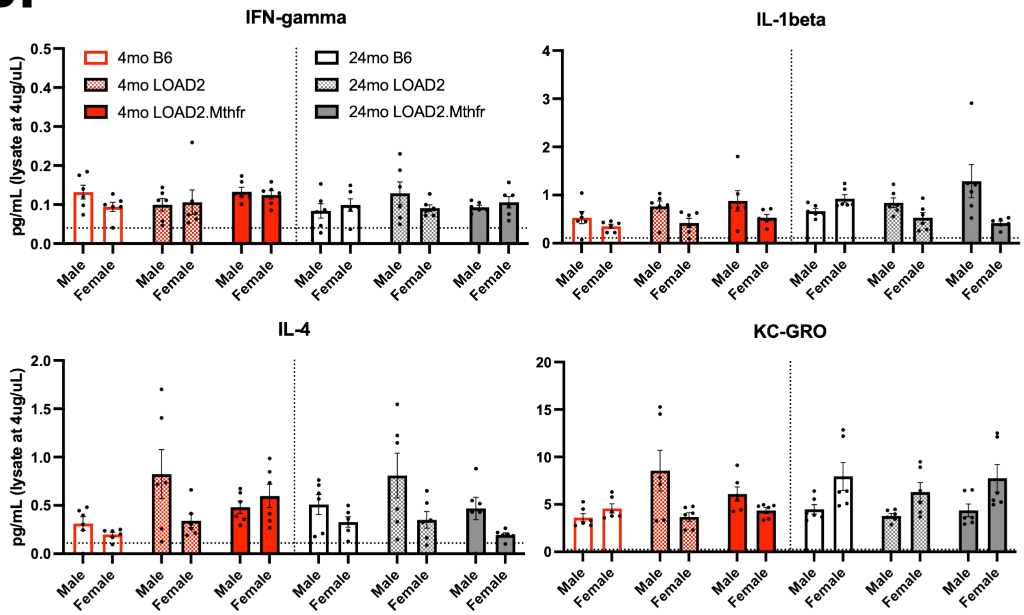

#### SUPP FIGURE 4

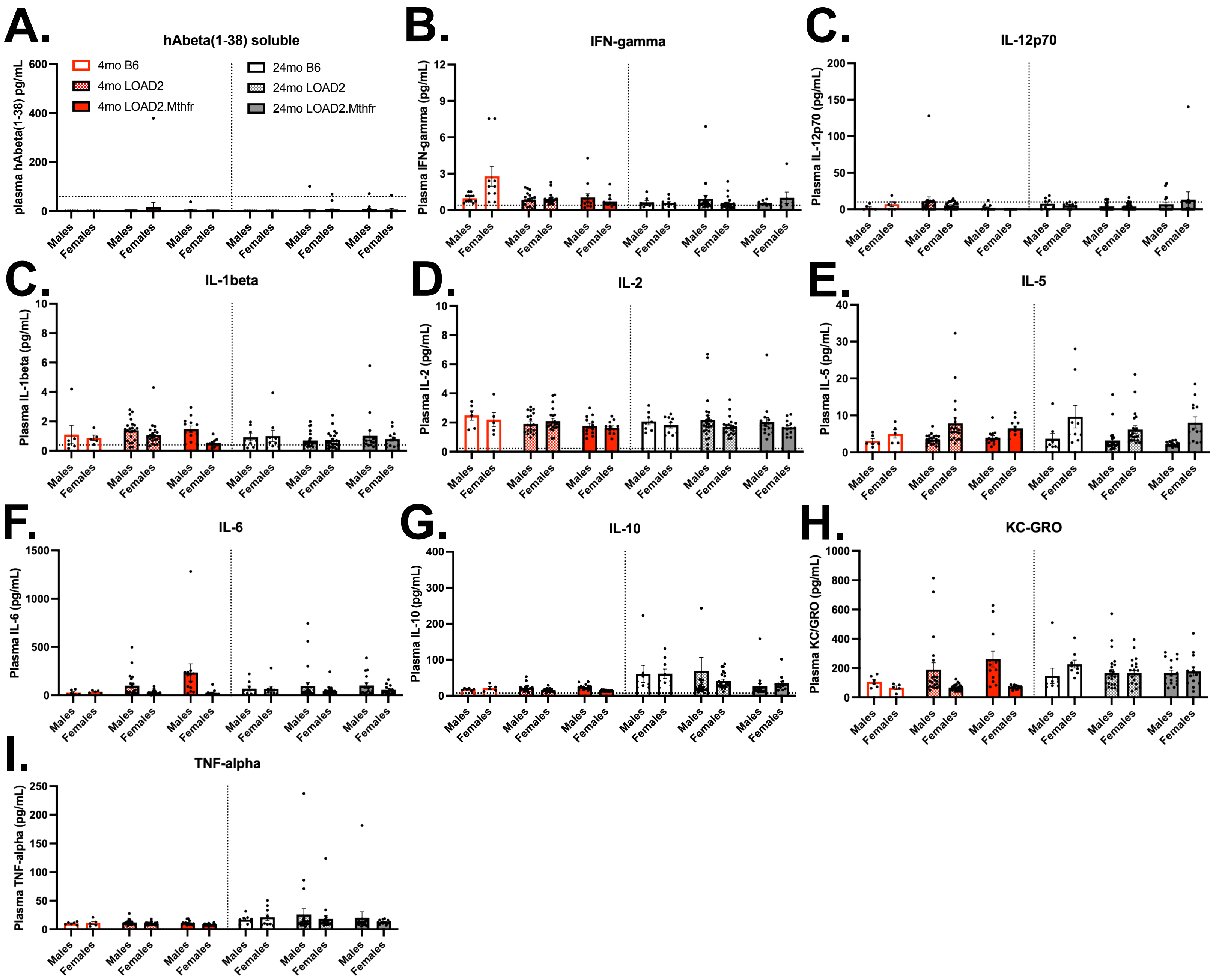

#### SUPP FIGURE 5

A.

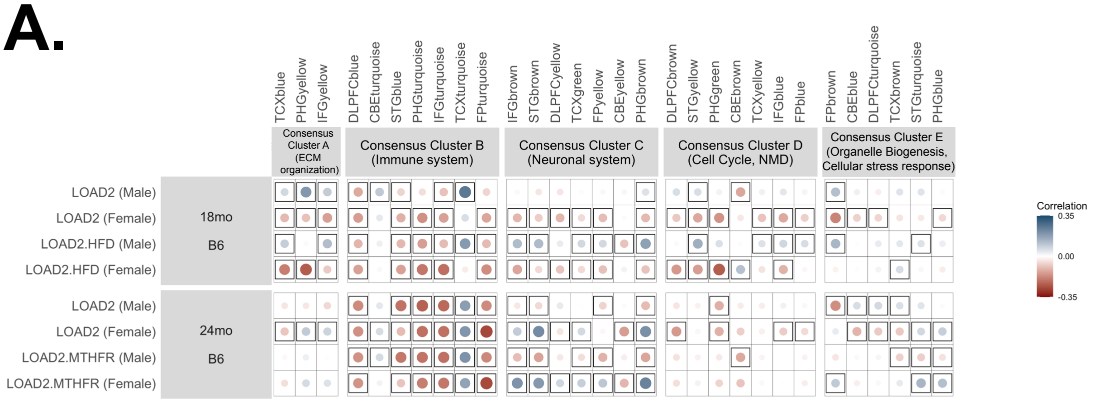


B.

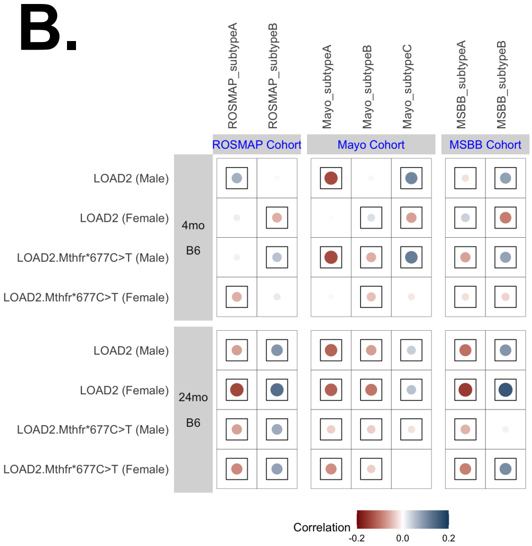


C.

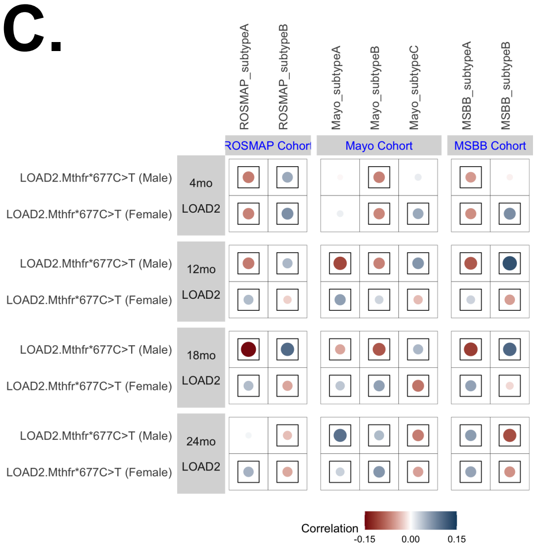
